# Robust switches in thalamic network activity require a timescale separation between sodium and T-type calcium channel activations

**DOI:** 10.1101/2020.10.08.327981

**Authors:** Kathleen Jacquerie, Guillaume Drion

## Abstract

Switches in brain states, synaptic plasticity and neuromodulation are fundamental processes in our brain that take place concomitantly across several spatial and timescales. All these processes target neuron intrinsic properties and connectivity to achieve specific physiological goals, raising the question of how they can operate without interfering with each other. Here, we highlight the central importance of a timescale separation in the activation of sodium and T-type calcium channels to sustain robust switches in brain states in thalamic neurons that are compatible with synaptic plasticity and neuromodulation. We quantify the role of this timescale separation by comparing the robustness of rhythms of six published conductance-based models at the cellular, circuit and network levels. We show that robust rhythm generation requires a T-type calcium channel activation whose kinetics are situated between sodium channel activation and T-type calcium channel inactivation in all models despite their quantitative differences.

## Introduction

Animal performance relies on their ability to quickly process, analyze and react to incoming events, as well as to learn from experience to constantly increase their knowledge about the environment. This information processing is shaped by fluctuations in rhythmic neuronal activities at the cellular and population levels, each defining *brain states* [Buzsáki (2009); Cannon et al. (2014)]. These activities are recognizable by spatiotemporal signatures of the mean-field activity of large neuronal populations. Switches between these brain states can be fast and localized, such as for example those observed in different brain areas prior to movement initiation [Kühn et al. (2004)], or global and long-lasting, such as those observed during the wake-sleep transition [McCormick, McGinley, and Salkoff (2015); McGinley et al. (2015)].

In the thalamus, cellular recordings reveal a firing pattern transition during wake-sleep cycles. The thalamic neurons switch from a regular spiking mode to a bursting mode [Guillery and Sherman (2002); McCormick and Bal (1997)]. This feature is not strictly restricted to sleep but can also appear during an awake and vigilant behavior [Cui and Shang (2011); Ding et al. (2016); Murray Sherman (2001); Ramcharan, Gnadt, and Sherman (2000); Sherman and Guillery (1996)]. At the population level, the change in cadence is remarkable; the mean-field activity rapidly switches from an active state to an oscillatory state [Zagha, Edward; Mc.Cormick (2014)].

The mechanisms governing rhythm fluctuations are poorly understood. This research problem is difficult to solve since it involves phenomena occurring at different scales; from molecular level to population level [X.-j. Wang (2010)]. The rhythms reflect a collective activity of neurons interconnected via synaptic connections. Besides, neuron dynamics are determined by a specific balance of ionic currents [Yu, Yarov-Yarov, Gutman, and Catterall (2005)]. These neuron intrinsic properties are controlled by neuroactive chemicals, called *neuromodulators*. Transition from sleep to wakefulness is associated with massive modifications in neuromodulators levels, such as serotonin, norepinephrine, dopamine and acetylcholine. These substances alter the behavior of thalamocortical neurons inducing shifts in population activity [Ding et al. (2016); McCormick and Bal (1997); McGinley et al. (2015); Murray Sherman (2001); Zagha, Edward; Mc.Cormick (2014)].

In parallel with these rhythm fluctuations, learning and memory are attributed to the ability of neurons to modify their connections with other cells based on experience, a property called *synaptic plasticity* [Abbott and Nelson (2000); Citri and Malenka (2008)]. Synaptic plasticity mechanisms often exploit the level of correlation in the activity of connected neurons (for example in spike-time dependent plasticity), and can therefore be affected by abrupt changes in neuronal excitability. Strong constraints are exerted on models of plasticity because neural circuits are adaptable to help animals to modify their behavior. At the same time, these circuits must be stable in spite of these changes. There is a balance between adaptability and stability [Abbott (2003); Turrigiano, Abbott, and Marder (1994)].

The coexistence of these mechanisms raises challenging questions: how can switches in brain states remain reliable despite constant rewiring of neuron connectivity? How is synaptic plasticity affected by switches in brain states? Indeed, little is known about how shifts in network rhythms influence synaptic plasticity, hence learning. One reason for this puzzle is that state-of-the-art computational models of switches in brain states have often focused on the role of connectivity changes in network rhythm modulation [Bevan, Magill, Terman, Bolam, and Wilson (2002); Destexhe, Contreras, and Steriade (1998); Esser, Hill, and Tononi (2009); Krishnan et al. (2016)]. Such models are not appropriate to study the impact of transient network oscillations on synaptic plasticity and learning, since the rhythmic switch itself relies on a disruption of the connectivity established through learning.

Recent evidence suggests that control of rhythmic activity can happen *at the cellular level*. No matter how fast or long-lasting are the transitions, they are controlled without affecting neither synaptic strength nor the circuit interconnection topology [Marder and Bucher (2007)]. Such mechanisms have been studied in small circuits, prominent examples being the crustacean stomatogastric system [Destexhe and Marder (2004); Marder and Bucher (2007); Marder, O’Leary, and Shruti (2014); Prinz, Bucher, and Marder (2004); Schulz, Goaillard, and Marder (2006)], and the heart [Olypher and Calabrese (2007); Roffman, Norris, and Calabrese (2012)]. These circuits are ideal supports to better understand how alteration in excitability regulates behavioral states. These works demonstrate that neuromodulation generates highly stable outputs. It has also been extensively shown that several combinations of ionic channels on the membrane or synaptic connections lead to the desired outcome [Marder, Abbott, Turrigiano, Liu, and Golowasch (1996); Schulz et al. (2006)]. It enhances the idea that rhythms cannot rely on a precise synaptic weight configuration or precise tuning of intrinsic parameters.

In this line of work; we aim at highlighting a cellular property that is critical for the generation of switches in brain states compatible with neuromodulation, cellular heterogeneity, synaptic plasticity and independent from network topology. This cellular property relies on a timescale separation between sodium and T-type calcium channel activations, providing a source of both fast and slow positive feedback loops at the cellular level. Slow positive feedback is accessible to all neurons that embed a sufficient amount of slowly activating voltage-gated calcium channels or slowly inactivating potassium channels in their membrane: the positive feedback comes from the fact that a calcium channel activation (resp. potassium channel inactivation) further amplifies the depolarization that gave rise to it, and slow means that calcium channel activation (resp. potassium channel inactivation) is at least one timescale slower than sodium channel activation. But the ultraslow inactivation of T-type calcium channels makes the slow positive feedback tunable: its gain depends on neuron polarization level. The presence of a slow positive feedback at the cellular level endows the neuron with an excitability switch, especially for the transition between tonic mode to bursting mode [Franci, Drion, Seutin, and Sepulchre (2013)].

This present work studies the role played by the timescale separation between sodium and T-type calcium channel activations on the robustness of six published thalamic neuron models. The thalamic neurons raise interest due to their varying firing patterns and contribution to brain states. Two models among the analyzed in this paper neglect this timescale separation by designing T-type calcium channel activation as an instantaneous event, a simplification often encounters in neuronal modeling. Here, we show through several computational experiments that the compatibility between neuromodulation, synaptic plasticity and switches in brain states correlates with the presence of the *slow* T-type calcium channel activation. As soon as intrinsic parameters and synaptic weights are affected respectively by neuromodulation and synaptic plasticity, the two models that speed up the calcium activation kinetics experience a drastic drop in their switching capabilities. However, restoring the slow activation of T-type calcium channels in these two models improves the robustness of their rhythmic activity.

To further quantify the importance of a timescale separation between sodium and T-type calcium channel activations, we vary T-type calcium channel activation kinetics in all models, ranging from the fast timescale of sodium channel activation to the ultraslow timescale of T-type calcium channel inactivation, and we test its robustness at the circuit level and at the population level. Our computational experiments confirm that the robustness of rhythmic activity is achieved when T-type calcium channel activation is an order of magnitude slower than sodium channel activation. This was observed in all models despite their quantitative differences. Our results thus highlight the importance of respecting the physiological timescale separation between sodium and T-type calcium channel activations to guarantee compatibility between neuromodulation, synaptic plasticity, cellular heterogeneity, adaptable connectivity and switches in neuronal rhythms.

## Results

### Robust vs. fragile firing pattern transition at the single-cell level

Throughout this paper, we compare six well-established conductance-based models of thalamic neurons [Drion, Dethier, Franci, and Sepulchre (2018)] (model 1), [Destexhe, Contreras, Steriade, Sejnowski, and Huguenard (1996)] (model 2), [Destexhe, Neubig, Ulrich, and Huguenard (1998)] (model 3), [Huguenard and McCormick (1992); McCormick and Huguenard (1992)] (model 4), [X. J. Wang (1994)] (model 5) and [Rush and Rinzel (1994)] (model 6). All these models include at least a sodium current, *I_Na_*, a potassium current, *I_K_*, T-type calcium, *I_CaT_* and a leak current, *I_leak_* (see SI for more details). Each of these models is conceived to reproduce the different firing patterns observed in a thalamic neuron (a depolarized tonic mode, a hyperpolarized bursting mode, rebound bursting, etc.) and the switch between them [Castro-Alamancos (2004); Guillery and Sherman (2002); McCormick and Bal (1997); Murray Sherman (2001)]. The firing mode is controlled by the external current. A depolarizing current drives the neuron model in tonic mode. If it is followed by an hyperpolarizing current, it switches the neuron model from a regular spiking mode to a bursting mode; a transition called hyperpolarized-induced bursting (Figure 1A) [McCormick and Bal (1997)].

**Figure 1:**
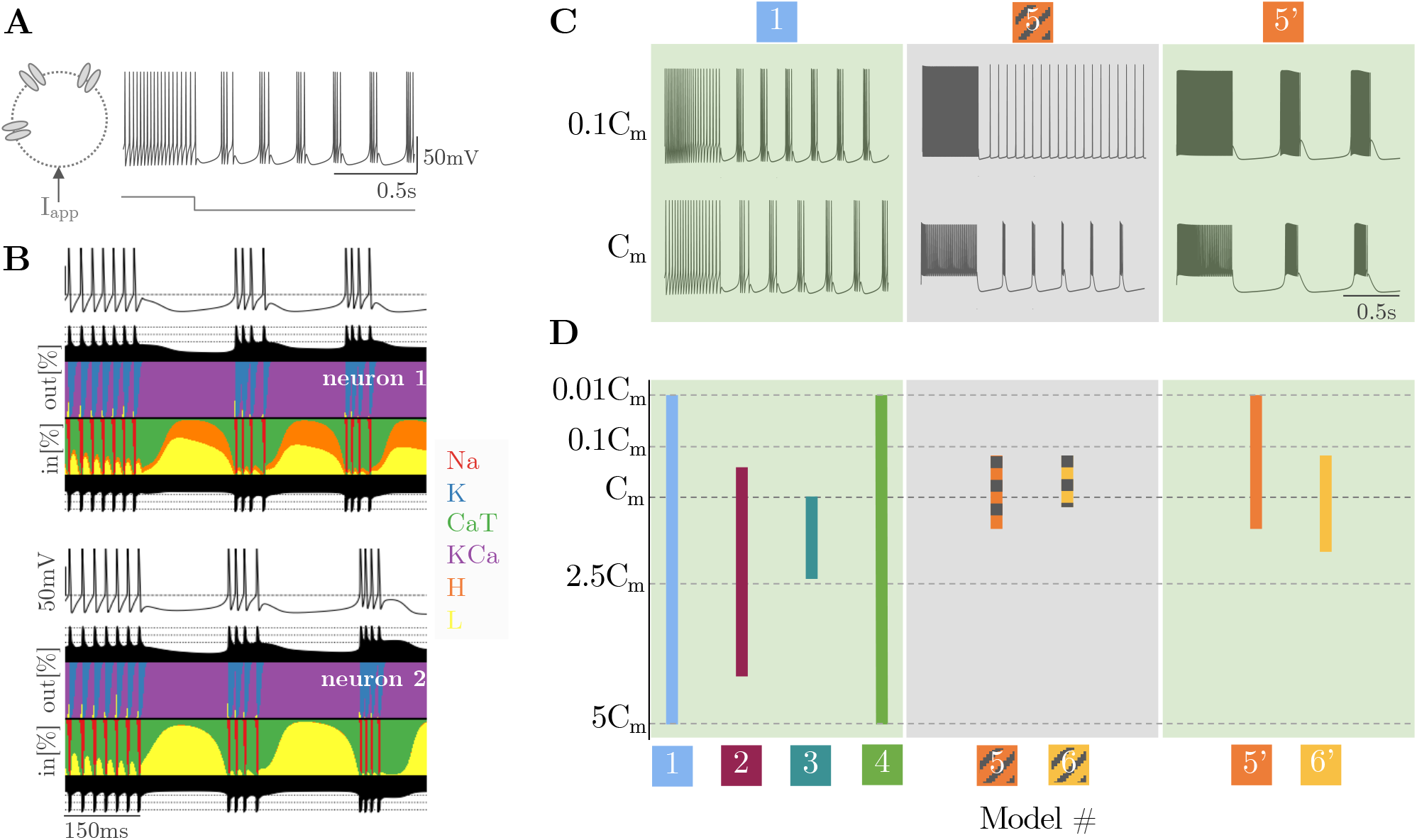
Models including slow activation of T-type calcium channels are robust to neuron variability such as change in cell size or in ion channel densities. **A.** Typical response of a model neuron when a hyperpolarized current is applied, called hyperpolarized-induced bursting (HIB). The cell switches from a regular tonic mode to a bursting mode (recording from model 1). **B.** Currentscape for a neuron model with two different sets of maximal intrinsic conductances. After 150ms, the two neurons are hyperpolarized. The membrane voltage time-course shows an HIB for both neurons. Under the membrane-time curve, the black surface shows respectively on the top and bottom total inward outward currents on a logarithmic scale (see [Alonso and Marder (2019)]). Between the black surfaces, each color curve reveals the contribution of one particular ionic current as the percentage of the total current during the simulation (recording from model 1). Both neurons display the same firing pattern but this outcome is achieved by different combinations of ion channels densities. **C** Only green models (models 1,2,3,4, 5’ and 6’) are robust to a uniform scaling of all the maximal conductances, modeled by a change in membrane capacitance (*C_m_*). The neuron model is able to switch from tonic to burst even though the capacitance membrane is divided by ten (left panel, recording from model 1). Gray models (models 5 and 6) are fragile to this parameter alteration. They lose the ability to switch (center panel, recording from model 5). Restoring the slow CaT channel activation is enough to recover the robustness (right panel, recording from model 5’). **D** Quantitative analysis of neuron model robustness to change in membrane capacitance. Each model is launched for a capacitance value varying from a hundredth to five times its nominal value. The parameter step is equal to 0.01*C_m_* (resp. 0.1*C_m_*) for simulations generated from 0.01 *C_m_* to 0.1*C_m_* (resp. from 0.1 *C_m_* to 5*C_m_*). Green models cover larger parameter range. By contrast, gray models lacking of slow CaT channel activation are fragile to membrane capacitance deviation. Replacing the instantaneous CaT channel activation by a slow activation provides the slow positive feedback mechanism to models 5 and 6. It turns these fragile models into more robust models. Hence, models 5’ and 6’ support larger variation of *C_m_*.

Experimental and computational studies have shown that a similar behavior or a similar firing pattern can emerge from neurons or circuits having very distinct intrinsic parameters [Alonso and Marder (2019); Goldman, Golowasch, Marder, and Abbott (2000); Marder and Prinz (2002)]. Figure 1B illustrates the membrane voltage time-course of a thalamic neuron model for two different parameter sets under the control of an external transient hyperpolarizing current (see black curves). The firing pattern is similar in both parameter sets and shows the typical switch occurring in thalamic neurons. However, the corresponding currentscapes reveal a variability in the contributions of the different ionic currents [Alonso and Marder (2019)]. A H-current is involved in the first neuron (top currentscape) while it is almost absent in the second neuron (bottom currentscape). It shows that intrinsic currents arrange themselves to maintain a given behavior. This simple experiment motivates the rest of this work; a computational model must be able to reproduce a desired outcome for a broad range of intrinsic parameters as it happens in biology.

Here, we study the robustness of conductance-based models to parameter variations with a special focus on the dynamics of the voltage-gated T-type calcium channel activation. The first four models (models 1 to 4, called *green* models) incorporate a *slow* activation of the T-type calcium (CaT) channels while the last two models (models 5 and 6, called *gray* models) fix the activation as an instantaneous event. This simplification is often encountered in neuronal modeling [Amarillo, Mato, and Nadal (2015); Golomb, Wang, and Rinzel (1994); Kubota and Rubin (2011); Pospischil et al. (2008); Rubin and Terman (2004); Rush and Rinzel (1994); Smith, Cox, Sherman, and Rinzel (2000); X. J. Wang (1994)]. Indeed, it removes one differential equation, which decreases the simulation time - a computation intake that is non-negligible as soon as one moves toward network simulations.

First, we investigate the impact of the CaT channel activation *dynamics* in the model robustness at the single cell level. To do so, we simply test the ability to reproduce the switch from tonic to burst by changing a single parameter in the model, namely, the capacitance membrane *C_m_*. An alteration in this parameter substitutes a change in cell size or a uniform scaling of all maximal ionic conductances [Franci, Drion, and Sepulchre (2018); O’Leary, Williams, Franci, and Marder (2014)]. Figure 1C reveals the striking consequence for the model robustness when the capacitance is scaled by a factor 1/10. Green models (models 1 to 4, left panel) including the slow activation of T-type calcium channel are able to reproduce the hyperpolarized-induced bursting while gray models that assume this activation is instantaneous are fragile. Indeed, models 5 and 6 (center panel) loose the ability to switch from tonic to burst. To make the comparison as fair as possible, we have restored the slow dynamics of the CaT channel activation in models 5 and 6. We reconstruct a differential equation for the activation variable whose time constant is voltage-dependent. Models 5*’* and 6*’* are their respective modified versions (see SI for details). This only computational change is sufficient to recover the switch from tonic to burst when the capacitance membrane is divided by 10 (Figure 1C, right panel).

To be more quantitative, this computational experiment is reproduced for membrane capacitance values scaled from a hundredth to five times its nominal value (see methods for details). Figure 1D reveals that green models are able to switch for a large range of capacitance values (see left panel). Gray models are fragile and cover a tiny range around the nominal value (*C_m_*) (center panel). Reinstating the slow dynamics of this channel bounces back the robustness as shown by the increase of the orange and yellow bars (right panel). Models 5*’* and 6*’* are able to generate the firing pattern switch typical in thalamic cells given capacitance values for which gray models are not able.

### Slow T-type calcium channel activation makes an isolated excitatory-inhibitory circuit robust to neuromodulation and synaptic plasticity

To extend our results obtained at the single-cell level, we move to the circuit level. We build an isolated excitatory-inhibitory circuit of two neurons. These neurons are connected through AMPA, GABA_A_ and GABA_B_ connections to model the asymmetric coupling between a subpopulation of excitatory (E) cells and a subpopulation of inhibitory (I) cells [Guillery and Sherman (2002); McCormick et al. (2015)]. This topology is a typical configuration in the thalamus [McCormick and Bal (1997); Sherman and Guillery (1996)]. The E-I circuit and its expected rhythmic network activity are illustrated in Figure 2A (left and center panels). It is controlled by an external current exerted on the inhibitory cell. Initially depolarized, the I-cell exhibits a tonic mode. The E-cell remains silent. As soon as the external current hyperpolarizes the I-cell; it deinactivates the T-type calcium channels, leading to a bursting mode. Then, thanks to the reciprocal connections, the circuit switches to a synchronous burst called the oscillatory mode.

**Figure 2:**
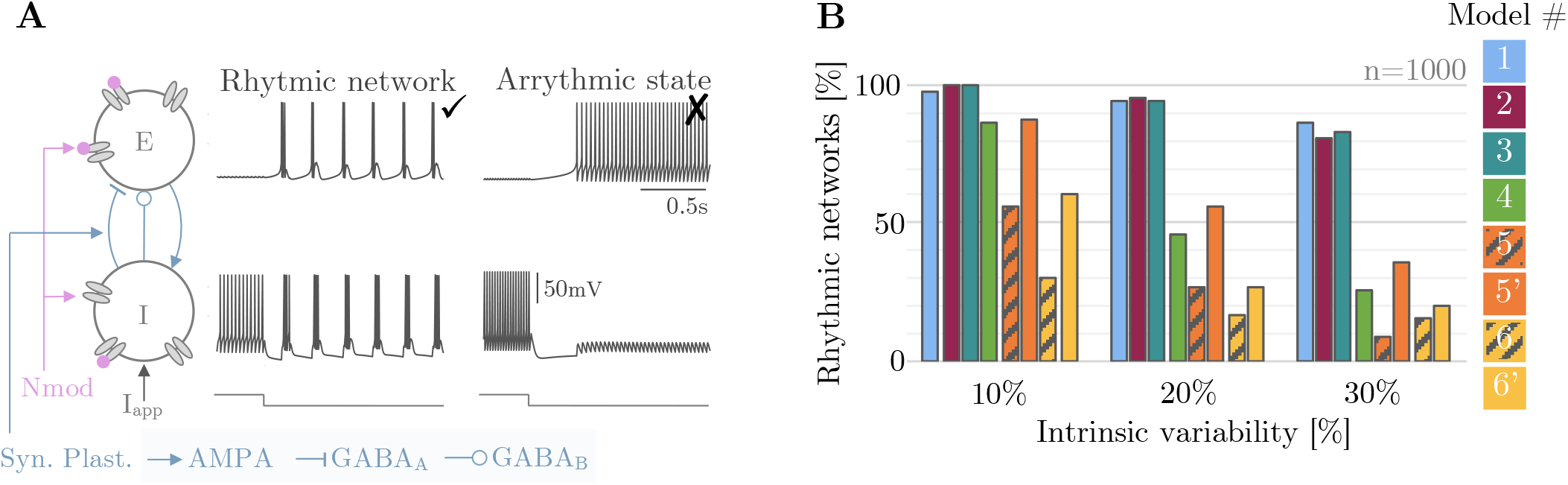
Models integrating slow activation of CaT channels generate robust network activities in presence of neuromodulation and synaptic plasticity. **A.** Two interconnected neurons (one excitatory neuron E and one inhibitory neuron I) under the control of an external current (I_app_), affected by neuromodulation (Nmod, in pink, represented by spheres) and synaptic plasticity (Syn. Plast., in grayish blue). The external current, initially depolarized, transiently hyperpolarizes the inhibitory cell and switches the rhythm of the circuit. The left traces shown that the initially silent excitatory neuron and the spiking inhibitory neuron start to synchronize their bursting phase. By contrast, the right traces show an example of arrhythmic circuits. **B.** Proportion in percentage of rhythmic networks observed in different neuron models as intrinsic variability increases. For each model, one thousand 2-cell circuits are generated with random ionic conductances varying from 10, 20 and 30% from their nominal values - mimicking the effect of neuromodulation, and synaptic conductances, randomly picked in a uniform distribution 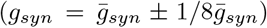 mimicking the effect of synaptic plasticity. For an intrinsic variability equals to 10%, models with slow CaT channel activation (models 1 to 4) show that more than 800 circuits switch from tonic to synchronous bursting mode while models with instantaneous CaT channel activation (models 5 and 6) are fragile to parameter variation. Restoring the slow timescale of the CaT channel activation strongly improves the model robustness (models 5’and 6’). For higher intrinsic variability(20% and 30%), models 1 to 3 maintain their ability to generate rhythmic networks. Model 4 is weaker due to its high number of ionic channels. For gray models (models 5 and 6), most of the thousand random circuits cannot switch whereas turning the instantaneous activation into a slow activation reveals a rise in robustness (models 5’ and 6’).

With a simple computational experiment, we study the robustness of these network switches to changes in neuron intrinsic properties, mimicking the effect of neuromodulation (modeled by maximal conductances), and changes in synaptic weights, mimicking the effect of synaptic plasticity (modeled by the synaptic conductances). For each model, we start from an E-I circuit capable of generating a switch. Then, one thousand 2-cell circuits are simulated by generating different parameter sets for maximal intrinsic conductances and synaptic weights. The maximal conductances vary within an interval of 10, 20 and 30 % around their nominal values and the synaptic weights vary in a fixed range (see methods for details). The percentage among the thousand 2-cell circuits that have performed the rhythmic transition is evaluated for the three intervals of variability in each conductance-based model. A rhythmic network is defined according to the firing pattern evolution shown in Figure 2A (center) while other activities are classified as an arrhythmic state, for example the pattern in Figure 2A (right). For a small intrinsic variability (10%), Figure 2B shows that models 1 to 4 are robust to neuromodulation and synaptic plasticity. More than 800 sets of parameters allow the circuit to switch. The absence of slow positive feedback in models 5 and 6 has a dramatic consequence on the model robustness. One every two parameter sets in model 5 cannot reproduce the typical thalamic activity transition. Model 6 is even more fragile. However, restoring the dynamical cellular property significantly improves the robustness as shown in models 5’ and 6’.

For larger intrinsic variability (20% and 30%), models 1 to 3 maintain their capabilities to switch for a broad range of intrinsic and synaptic parameter ranges. Model 4 can be considered apart. It is shrinking as the variability increases. This decrease in performance is likely related to its high number of conductances (about twice as much as the other models). Indeed, we are exploring a 14-dimension space since this model has 11 intrinsic conductances and 3 synaptic conductances. The parameter exploration for the other models occurs in a 7 to 9 dimension space reducing the model complexity. Models 5 and 6 have a similar number of intrinsic conductances as models 1-3 but they are extremely fragile to parameter changes. They are almost unable to perform rhythmic transition. The rhythm in an E-I circuit requires a precise tuning of intrinsic and synaptic parameters for models lacking of the slow kinetics of CaT channel activation. This computational choice makes the model incompatible with neuromodulation and synaptic plasticity. Once again, their modified versions embedding the slow positive feedback (correlated to the slow activation of T-type calcium channels) have a better response to parameter perturbations.

### A timescale separation between sodium and T-type calcium channel activations ensures compatibility between circuit switch, neuromodulation and synaptic plasticity

So far, we have shown that modeling the T-type calcium channel activation with a slow kinetics drastically enhances the robustness of rhythmic switches at the cellular, and circuit levels. But one question remains: what does *slow* mean, and how tuned does the activation kinetics need to be to achieve robustness? Indeed, their exists different subtypes of T-type calcium channels whose activation kinetics can greatly differ [Hille (2001); Perez-Reyes (2003)].

To answer this question, we explore the impact of incrementally varying T-type calcium channel activation kinetics in a similar computational experiment as done in Figure 2. We focus on green models (models 1, 2, 3, 4, 5’ and 6’) that describe the opening of the T-type calcium channel with a first order differential equation 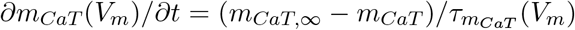. The variable 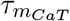 is the voltage-dependent activation time constant and characterizes the dynamics of channel opening. We start from an isolated 2-cell E-I circuit connected via AMPA, GABA_A_ and GABA_B_ synapses. The circuit is able to switch from tonic to burst at the nominal parameter values. Then, we add neuromodulatory effect - by varying intrinsic parameters, and synaptic plasticity - by changing extrinsic parameters. We build four hundred 2-cell circuits whose maximal ionic conductances and synaptic conductances are randomly picked in an interval of 20% around their basal values. Here, the novelty is to play with the time constant of CaT channel activation 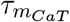 of the two neurons (see Figure 3A).

**Figure 3:**
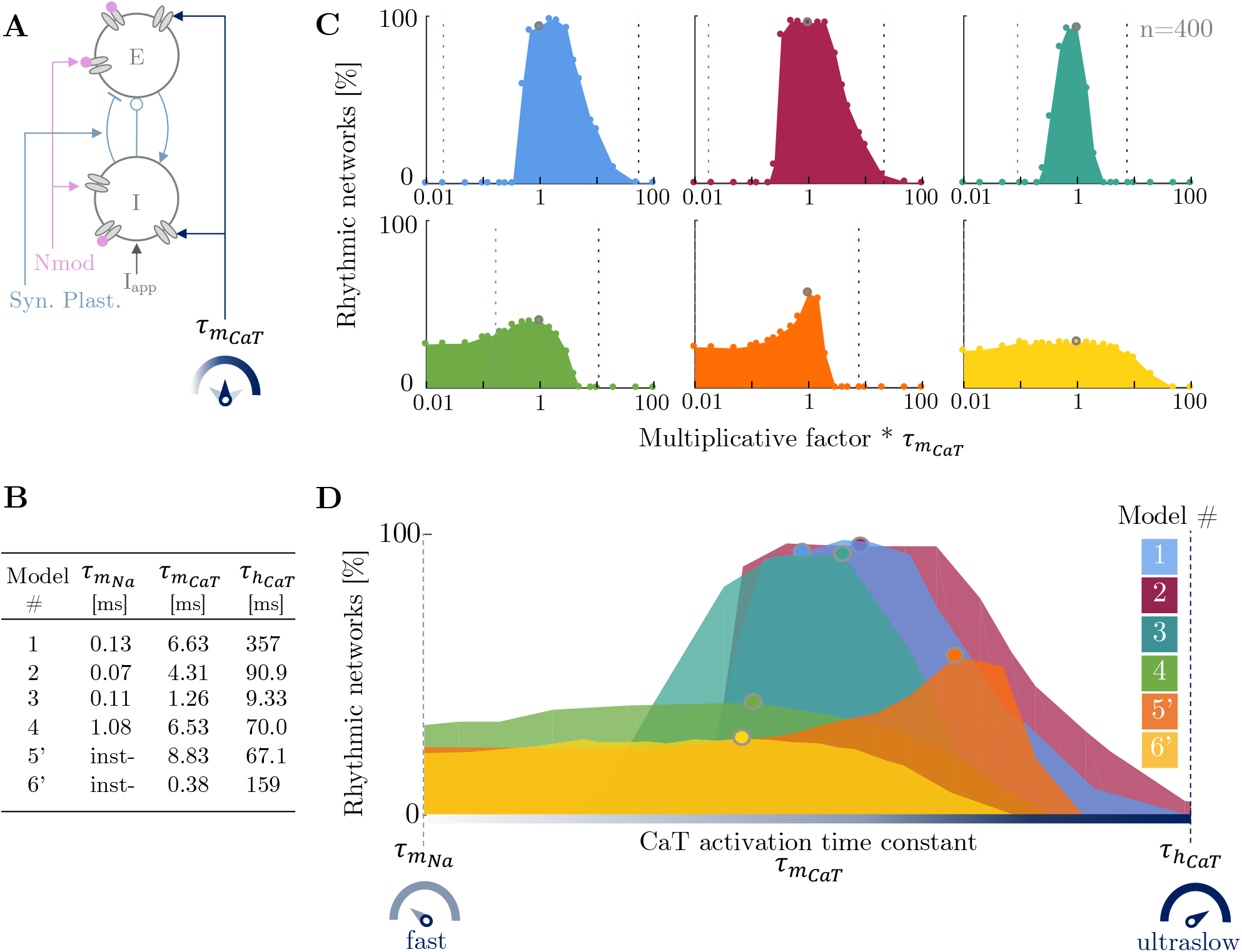
The physiological *slow* timescale of CaT channel activation guarantees compatibility between circuit switch, neuromodulation and synaptic plasticity. **A.** For each model, 400 E-I 2-cell circuits are built. In each circuit, the intrinsic and synaptic conductances are randomly picked in a uniform range of *±* 20 % around their nominal values to mimic neuromodulation and synaptic plasticity. These same 400 circuits are then simulated for a varying CaT channel activation time constant 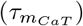. **B.** Comparison table between the three time constants associated with sodium channel activation 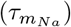, CaT channel activation 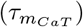 and CaT channel inactivation 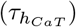 evaluated at their threshold value (see methods for details). For models 5’ and 6’, “inst-” means the sodium channel activation is modeled as an instantaneous event. There is not differential equation describing its kinetics, consequently, there is no voltage-dependent time constant. The numbers greatly widely vary between each model. It confirms the quantitative differences between them. Effect of a varying CaT current activation time constant on the switching capability in a 2-cell circuit for models 1, 2, 3, 4, 5’ and 6’. The y-axis quantifies the percent of rhythmic networks among the 400 simulated random circuits under different values of 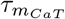. The parameter 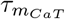 is scaled by a multiplicative factor varying from 0.01 to 100. The smallest the multiplicative factor, the fastest the CaT channel activation. The performance associated with the nominal CaT activation time constant is depicted with a gray circle while the results associated with the scaled time constants are shown with full circles. The sodium channel activation (resp. CaT channel inactivation) time constant is marked with the left (resp. right) dashed vertical line. For models 5’ and 6’, the Na current activation is instantaneous. For model 6’, the CaT channel inactivation is greater than one hundred times its activation. For each model, the circuit robustness shrinks as soon as the channel is opening too fast or too slow. **D.** To compare models within each other, the x-axis is a *normalized* logarithmic scale such as: 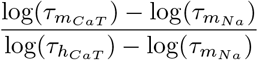. Each model is superposed between a window bounded on *τ τ* the left (resp. right) by the sodium channel activation (resp. CaT channel inactivation) time constant. The curve profile of all models confirms that the CaT activation time constant must remain confined in the *slow* timescale in order to guarantee model robustness to neuromodulation and synaptic plasticity. Decreasing the 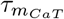, *i.e.* leading to a faster calcium activation makes the model fragile. Similarly, when 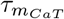 is increased, *i.e.* driving the time constant to the ultraslow timescale, none circuit switches from tonic to burst.

Figure 3B is a comparative table between time constants associated to sodium channel activation 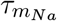, CaT channel activation 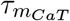 and inactivation 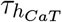 evaluated at their threshold voltage (see methods for detail). It points out the quantitative differences between models. Despite these differences, we are questioning on the choice made for 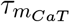 with respect to 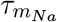 and 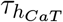. To do so, the time constant 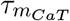 is scaled by several multiplicative factors from 0.01 to 100 times its nominal value. The smallest the coefficient, the fastest the CaT channel activation. For each scaled CaT time constant, we test the model capability to switch from tonic to burst when the I-cell is hyperpolarized. Among the 400 tested circuits, the percentage of rhythmic circuits is placed on the y-axis (see Figure 3C). The x-axis is on a logarithmic graduation. When the multiplicative factor is equal to one (marked by the gray circle), it indicates the CaT channel activation time constant initially designed for each model. The sodium channel activation time constant 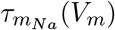 and the CaT channel inactivation time constant 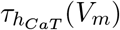 are also drawn for each model in dashed lines (respectively 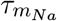 on the left and 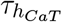 on the right). They are evaluated at their threshold voltage (see methods for details). They respectively indicate the *fast* and the *ultraslow* timescale. For models 5’ and 6’, the sodium current activation is instantaneous. For model 6’, the CaT channel inactivation is greater than 100 times its activation, it does not appear on the graph.

The outcome of this computational experiment is compelling. Figure 3C reveals that models are robust to neuromodulation and synaptic plasticity when the timescale of the CaT channel is situated in a slow range. The meaning of slow stands by itself; it is bounded between the fast kinetics of the sodium channel activation and the ultraslow kinetics of CaT channel inactivation. For each model, the peak of robustness lies between these two timescales. The bump-shaped surface shows a relatively large width. It points out that the activation kinetics do not need to be perfectly equal to one specific value between the fast and ultraslow regions. But these kinetics just need to be included within the slow timescale range. However, as soon as the kinetics moves too far from this interval, the robustness loss is abrupt.

To go further in the analysis, the six models are superposed on each other by normalizing the logarithmic x-axis on Figure 3D. The left (resp. right) boundary is the time constant of the sodium channel activation (resp. CaT channel inactivation); namely, the fast and the ultraslow timescales. The number of rhythmic circuits is enhanced when the CaT activation occurs at a timescale slower than the sodium activation and faster than the CaT inactivation as highlighted by the bump-shaped surface. Modeling the CaT channel opening at a slow timescale guarantees the compatibility between neuromodulation, synaptic plasticity and switches in brain states. Indeed, the compatibility relies on the presence of a *slow positive feedback* at the cellular level, as mentioned above.

Model 4 maintains a steady robustness even if the CaT channel activation is accelerated. This can be explained by the presence of another source of slow positive feedback such as a slowly activating L-type calcium channels (see SI for more details about model 4). Models 5’ and 6’ display a modest robustness for a fast opening of the CaT channel. These models were initially designed to operate for an instantaneous activation. However, the favorable operating point is preferably at a slow timescale.

If the kinetics of the CaT channel opening slows down too much (meaning we are moving to the right on the x-axis), it reaches the same timescale as its inactivation. In other words, the activation gate opens while the inactivation gate closes leading to a zero flux of calcium ions. The kinetics of CaT channel activation must be slow but not too slow.

### A timescale separation between sodium and T-type calcium channel activations promotes robustness of network states in large heterogeneous populations

From an isolated E-I circuit of two neurons, we build a larger network whose topology is emblematic of the thalamus, illustrating the interaction between relay neurons and the reticular nucleus. This population interaction is involved in state regulation such as the transition from wakefulness to sleep [McCormick and Bal (1997); Murray Sherman (2001)]. We are repeating the two previous computational experiments performed at the circuit level, now on a neuronal population. To do so, we start with a 200-cell network where the population of 100 excitatory neurons is identical to the population of 100 inhibitory neurons. We neglect intra-population interaction, and we assume all-to-all connectivity between the two populations. The E-cells project AMPA synapses to all the I-cells and conversely, the I-cells are connected to the E-cells via GABA_A_ and GABA_B_ synapses. All the synaptic weights linking the neurons together are identical. An external current is exerted on the inhibitory cells (see Figure 4A). This hyperpolarizing current causes a cellular switch and drives the neurons in a synchronous bursting mode as previously shown with the isolated E-I circuit in Figure 2A [Drion et al. (2018)]. This change in cellular firing pattern is translated by an oscillatory behavior at the network level. This oscillatory state can be visualized by computing the local field potential (LFP) of the neuron population. LFP is measured as the sum of synaptic activity in a neuronal population (see Figure 4B, top curves). When the current hyperpolarizes the I-cells, the synaptic current is remarkably modified and reveals a stronger activity. The spectrogram of the LFP shows that the transient hyperpolarizing current turns on the mean-field rhythmic activity marked by a strong power LFP frequency band. For each model, the homogeneous network is able to switch from an active state to an oscillatory state (see Figure 4B).

**Figure 4:**
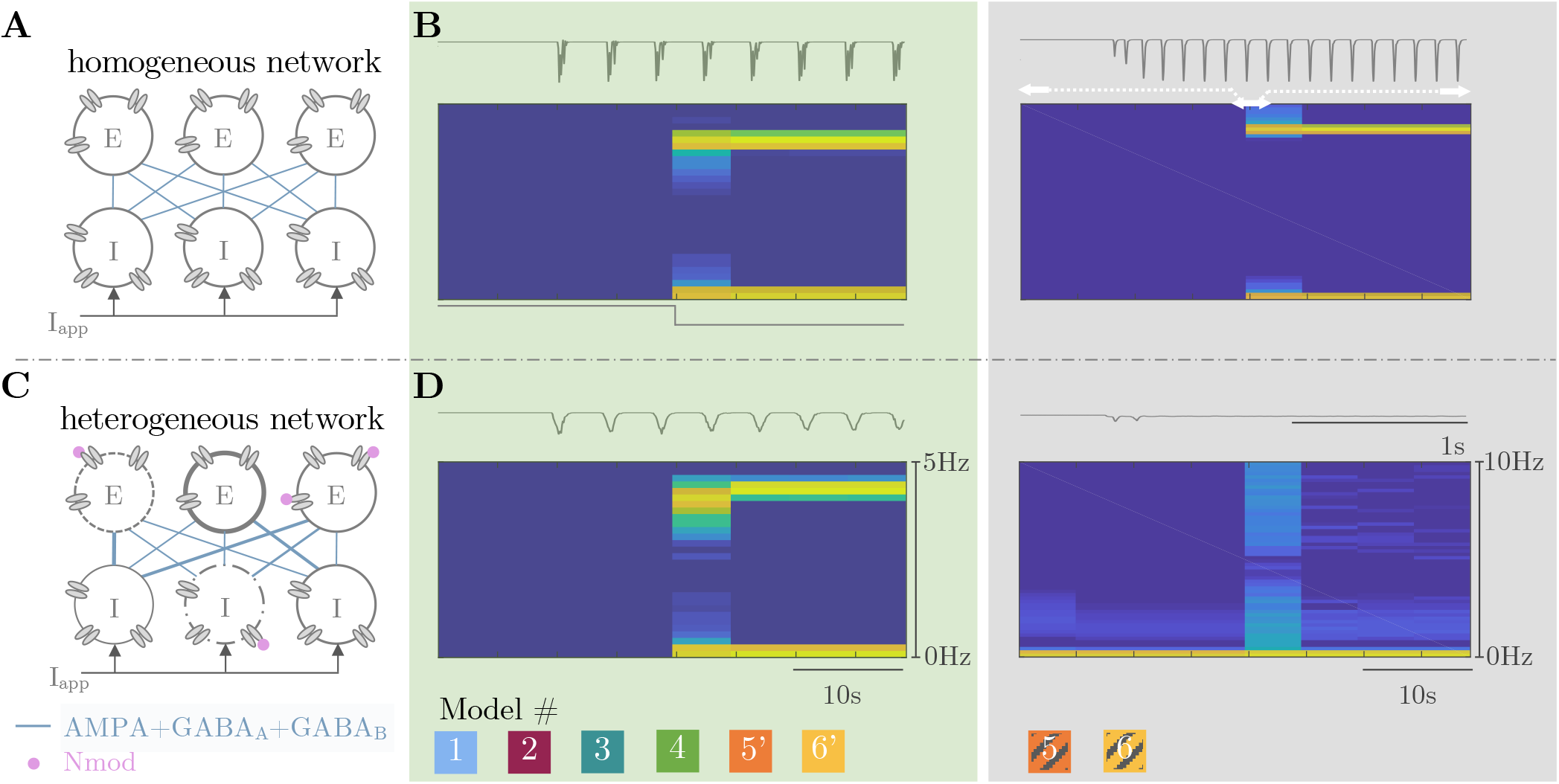
Models embedding a slow activation of CaT channels generate robust network switch independent of the network topology and the neuron heterogeneity. **A.** A 200-cell network is built with the same neuron model; 100 excitatory cells connected via the AMPA synapses to 100 inhibitory cells projecting back GABA_A_ and GABA_B_ synapses. The network is homogeneous. All neurons have the same channel densities and the same synaptic weights. **B.** Time-course (top) and spectrogram (bottom) of the local field potentials (LFPs) of the inhibitory neuron population for the homogeneous network. The external current transiently hyperpolarizes the inhibitory neurons. It turns on the mean-field rhythm activity of the population depicted by a stronger synaptic activity on the LFP time course and by the transient high power LFP frequency band on the spectrogram. **C.** A heterogeneous 200-cell network is built to take into account neuromodulation, cell variability, more representative topology and synaptic plasticity. Each ionic conductance is randomly picked in a given range around its nominal value. Each synaptic conductance is selected in a uniform distribution 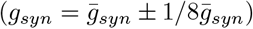. **D.** (left panel) Only green models (models 1,2,3,4 5’ and 6’) display the switch in the mean field rhythm of the population marked with an oscillatory LFP time-course (top) and a significant power band in the spectrogram (bottom). (right panel) Gray models (models 5 and 6) that lack slow CaT channel activation are fragile to the network topology and the heterogeneity. Switch in population rhythm is recovered when the slow regenerativity is restored (models 5’ and 6’).

However, a perfect network with identical neurons and identical synaptic weights is not consistent with reality. Each neuron differs from its neighbor with different intrinsic parameters such as the cell size or the densities of ionic channels. In addition, connections between neurons are neither identical nor static. Therefore, we explore the model ability to maintain switch in brain states in presence of cellular heterogeneity and its independence on network topology [Drion et al. (2018); Drion, Franci, and Sepulchre (2019)]. To do so, we build a 200-cell network where each neuron is different and the connectivity is uneven (see Figure 4C). The intrinsic and synaptic parameters are randomly picked in a given interval (see methods for details). Figure 4D shows the astonishing contrast between the stability of green models including the slow activation (see left panel) and the fragility of gray models lacking of this property (see right panel). Models 5 and 6 are able to switch from an active state to an oscillatory when the network is homogeneous and the connectivity is perfectly balanced. However, as soon as the network is changed into a more realistic configuration, these models cannot preserve switches in brain states as shown with the flat LFP curve or the absence of a marked power band in the spectrogram. Models 1 to 3 show a marked power band in their spectrogram when intrinsic parameters vary in an interval of 20% around their nominal values. Once again, model 4 is less robust, certainly due to its high number of conductances. It continues to switch for an intrinsic variability of 10%. Model 5 (resp. model 6) does not tolerate a variability of 20% (resp. 5%). When the slow activation of T-type calcium channels is reestablished, models 5’ and 6’ switch from an active state to an oscillatory state at the same level of variability that the initial model was fragile. This modeling modification leads to a neuron model robust to cell variability and that does not rely on the network topology.

To go further, we explore once again the relevance of respecting physiological timescale separation in ionic current modeling but this time at the population level. To do so, we build a 200-cell network with 100 excitatory cells connected to 100 inhibitory cells via AMPA, GABA_A_ and GABA_B_ synapses. Models 1 to 4, 5’ and 6’ are switching the network state under a hyperpolarizing current for a homogeneous and a heterogeneous configuration, as shown in Figures 4 (green models). Here, we test which kinetics provides robustness to network heterogeneity. The CaT activation time constant 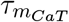 of the 200 neurons are scaled by several multiplicative factors ranging from an eighth to eight times its nominal value. In addition, cellular and synaptic variabilities are introduced by randomly picking the maximal intrinsic and extrinsic conductances of each neuron in a given interval around their nominal values following a uniform distribution (see Figure 5A). These heterogeneous networks are constructed with parameters varying in an interval whose width is ranging from 0 to 50% with a step of 5%. At each scaled 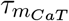, we test the maximal possible variability width at which the heterogeneous network is able to switch. For example, we build the 200-cell network whose maximal ionic and synaptic conductances are picked in a range of 20% around their initial values. Then, we simulate the network under the control of an hyperpolarizing current at different scaled 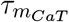. We finally check for each timescale if the given heterogeneous network is switching or not, by analyzing its LFP activity. The network is switching if the time course of neuron population LFP displays an oscillatory behavior in the hyperpolarized state and if its spectrogram is marked by a strong power band (see Figure 5B). We continue to increase the variability interval to quantify the correlation between the timescale and its robustness to cellular heterogeneity and uneven topology.

**Figure 5:**
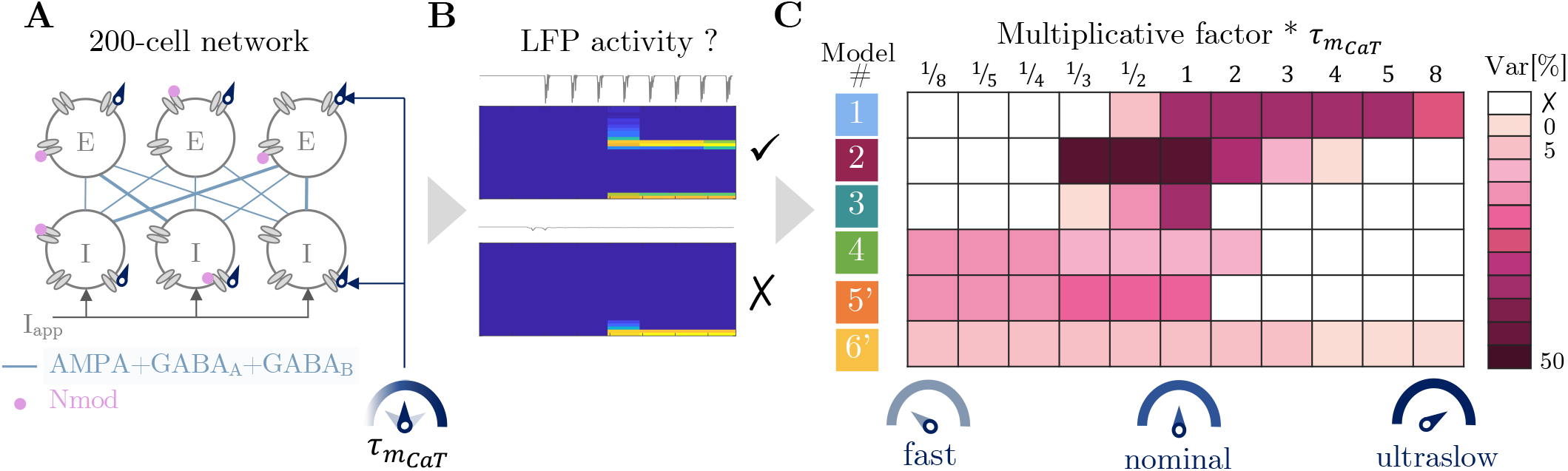
The physiological *slow* timescale of CaT channel activation yields to rhythm switches independent of the network topology and cell variability. **A.** A 200-cell network with 100 excitatory neurons and 100 inhibitory neurons connected via AMPA, GABA_A_ and GABA_B_ synapses under the control of an hyperpolarizing current. The intrinsic and extrinsic parameters are respectively affected by neuromodulation and synaptic plasticity; their values are randomly picked in a given interval. The CaT channel activation time constant 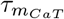 of each neuron is scaled by a multiplicative factor. **B.** At each scaled time constant, we check if the heterogeneous network is switching by analyzing the LFP activity. If the LFP timecourse presents a strong activity and its spectrogram shows a marked power band, the network displays an oscillatory state during the hyperpolarization state. **C.** The table summarizes the largest variability width at which the network presents a switch in its mean-field activity for several scaled time constants. Respecting the slow timescale of the CaT channel activation guarantees the switch in network rhythm compatible with variability in channel densities and synaptic weights. Driving the CaT channel activation to a fast or ultraslow timescale makes models more fragile to network topology and heterogeneity. Model 4 maintains its ability to switch from an active state to an oscillatory state even if the CaT activation time constant decreases. Its robustness comes from another source of slow positive feedback.

Figure 5C summarizes the model robustness at each scaled CaT time constant. It confirms our previous result; accelerating or decelerating the CaT channel opening makes the six models fragile to heterogeneity. In models 1 to 3, the best operating point to set the CaT time constant is confined between the fast timescale and the ultraslow timescale as shown by the darker zone. It enhances the model capability to switch in presence of cellular heterogeneity and synaptic plasticity. Model 4 is not really robust due to its high number of ionic currents. It is also assumed to embed another source of slow regenerativity helping him to operate at a faster timescale. As exhibited in Figure 3C for an isolated 2-cell circuit, models 5’ and 6’ also maintain a certain ability to switch even at a faster timescale because they were initially designed to operate at an instantaneous opening of CaT channels. As adapted versions of models designed to switch with an instantaneous T-type calcium activation, their robustness is lower than models 1 to 3 in all parameter ranges.

Overall, Figure 3 and Figure 5 reveal the importance of considering the physiological kinetics range of ion channel gating in computational models and especially the timescale separation between the different ionic currents. Getting rid of the slow dynamics of the T-type calcium channel activation disrupts the timescale separation with sodium channel activation. It removes an important biological property of this neuron type and thus, disturb its ability to change its firing pattern under a transient hyperpolarization in presence of neuromodulation and synaptic plasticity. This deficiency is transposed at the population level as shown by the inability to turn on the mean-field activity as soon as the cellular heterogeneity and unbalanced connectivity are increased.

## Discussion

### The physiological timescale separation between sodium and T-type calcium channel activations

A quotation from Bertil Hille’s book [Hille (2001)], “The time course of rapid activation and inactivation of T-type *I_Ca_* has been described by an *m*^3^*h* model like that for *I_Na_* (Coulter, Huguenard, and Prince (1989); Herrington and Lingle (1992)), but the derived time constants *τ_m_* and *τ_h_* are 20 to 50 times longer than those for *I_Na_* of an axon at the same temperature.”, highlights a physiological timescale separation between sodium and T-type calcium channel gating kinetics. T-type calcium channel presents of portfolio of activation kinetics depending on their isotype (i.e. Ca_*V*_ 3.1, Ca_*V*_ 3.2 or Ca_*V*_ 3.3), but all isotypes share the property of activating a timescale slower than sodium channels: the activation time constant ranges between 1ms and 50ms [Cain and Snutch (2010); Chemin et al. (2002); Choi, Yu, Lee, and Llinás (2015); Clapham and Garbers (2005); Frazier et al. (2001); Klöckner et al. (1999); Perez-Reyes (2003)]. By contrast, the activation time constant of sodium channel is lower than 1ms [Gilly, Gillette, and McFarlane (1997); Hille (2001); Hodgkin and Huxley (1952); Reckziegel, Beck, Schramm, Elger, and Urban (1998)]. As explicitly shown with these numbers, there is one order of magnitude between these two gating time constants.

### Modeling T-type calcium channel activation in conductance-based models

In a conductance-based model, the opening of an ion channel is represented by a voltage-dependent gating variable. This variable is described with a first order differential equation whose time constant is also voltage-dependent. In this paper, we investigated the kinetics chosen for the T-type calcium channel activation with respect to sodium channel activation. In other words, we compared these time constants with respect to each other.

There is a growing trend to design T-type calcium channel activation as the equivalent of a sodium channel activation. It often based on the fact that these two channels activate on a faster timescale than their own inactivation. For example in the published paper associated to model 5, it is said that “the activation variable [of T-type calcium channel] *s* is relatively fast and is replaced by its equilibrium function. (…) The activation kinetics [of the sodium current] being fast, the variable *m* is replaced by its equilibrium function (…) ” [X. J. Wang (1994)]. However, in the original version published three years before, the activation of T-type calcium channel was not at the same timescale as the sodium channel activation [X. J. Wang, Rinzel, and Rogawski (1991)]. In the same way for model 6, it is written “We employ a simplified version of the quantitative model originally formulated by Wang et al. (1991). Activation *s* is rapid, and inactivation *h* is relatively slow, (…) activation is assumed to be instantaneous. ” [Rush and Rinzel (1994)]. The similar simplification occurs for model 3 published in 1998: ten years later, the author mentioned “Note that the activation variable *s* is considered here at steady-state, because the activation in fast compared to inactivation. This T-current model was also used with an independent activation variable (Destexhe, Neubig, et al. (1998);Fig. 10), but produced very similar results as the model with activation at steady-state (not shown)” [Pospischil et al. (2008)]. Modeling the T-type calcium channel activation on the same timescale as the sodium channel activiation is a modeling simplification that does not only appears in models of thalamic neurons. For example in basal ganglia, subthalamic neurons are excitatory neurons projecting to inhibitory globus pallidus neurons. These neurons present a switch in firing activity that leads to different brain states, particularly relevant in movement generation and in Parkinson’s disease [Bevan et al. (2002); Brown and Williams (2005); Gatev, Darbin, and Wichmann (2006); Kühn et al. (2004); Swann et al. (2011); Uhlhaas and Singer (2006)]. Subthalamic neurons also embed T-type calcium channels. Once again, it is common to see that the activation of sodium and T-type calcium channels are modeled on the same timescale or even considered as instantaneous [Kubota and Rubin (2011); Rubin and Terman (2004); Terman, Rubin, Yew, and Wilson (2002)].

In both situations, these models are operational. It means that differentiating the sodium and T-type calcium channel activations is *not indispensable to reproduce* discharge modes. Those models are able to fire in tonic and burst. This timescale separation between the activation of these two different channels is neglected because it seems to be a computational detail. Figures 3 and 5 confirm that models initially considering a fast activation of T-type calcium channel can perform the desired activity. However, our results stress the crucial importance of this timescale separation for the *robustness* in network switches. The optimum operating timescale for T-type calcium channel activation is located between the fast activation of sodium channels and the ultraslow inactivation of T-type calcium channels.

As mentioned earlier, both channels have different electrophysiological properties. Assuming the T-type calcium channel activation as fast as sodium channel activation means that it is just a current summation. This has for consequence to drastically reduce the ability of generating a robust network activity as shown in our computational experiments.

### Compatibility between switches in brain states, synaptic plasticity and neuromodulation

Neuronal oscillations, synaptic plasticity and neuromodulation are hot topics in neuroscience since they are building blocks for information processing, learning, memory or adaptability. Studying the interaction between them requires computational models that generate *robust* activity even if the synaptic connectivity and the endogenous parameters are altered.

Altogether, this paper reveals that the *robustness* in the brain state switches in presence of cellular variability, heterogeneity at the network level is *correlated with the timescale separation* between sodium and T-type calcium channel activations. Despite the quantitative differences between the models, this timescale separation makes the rhythmic transition compatible with temporal variability and spatial heterogeneity induced by regulatory functions like neuromodulation, synaptic plasticity and homeostasis. In particular, triggering oscillations without any modification of synaptic connectivity makes the models well suited to study how a change in network activity can affect learning.

The mechanism highlighted in this paper can be exploited in other models than thalamic neuron models. It is can be extended to neurons that embeds slow-activating voltage-gated calcium channels or slow-inactivating potassium channels [Franci et al. (2013)]. The term slow confirms that the model should conserve the timescale distinction between the fast activation of sodium channels and the slow operating timescale of these two specified channels. This physiological timescale separation is imposed by ion channel dynamics. In conductance-based models, this corresponds to the time constant of the differential equation associated to the channel gating variable. It means that the time constant associated to sodium channel activation is on order of magnitude smaller than the time constant associated to the T-calcium channel activation.

Talking about positive feedback sources can be another approach to support the importance of differentiating ion channel kinetics. On one hand, sodium channel activation acts as a *source of fast positive feedback* because depolarizing the cell drives sodium ions in, making the cell even more depolarized and allowing more and more sodium ions to enter and so on. And the other hand, calcium channel activation operates in a similar way; it is also a source of positive feedback. However, there is a crucial dynamical difference between these two sources: they operates on two different timescales. If the two sources are acting on the same timescale, it is simply a sum of positive feedbacks. As shown by our experiments, it significantly decreases the robustness in presence of neuromodulation and synaptic plasticity. Each type of channel activation or inactivation that is a source of slow positive feedback, for example slow-activating voltage-gated calcium channels or slow-inactivating potassium channels, must be modeled an order of magnitude slower than the sodium channel activation.

Beyond the argument based on the time constant or the source of positive feedback, this time scale separation can be translated into dynamical analyses, in reduced models such as improved version of Fitzhugh-Nagumo model [Drion, Franci, Seutin, and Sepulchre (2012)] or integrate and fire models [Van Pottelbergh, Drion, and Sepulchre (2018)]. The secret is far from adding a complicated equation, on contrary, it simply relies on the timescale separation between sodium activation, a source of fast positive feedback and other currents that bring a source of slow positive feedback.

## Acknowledgment

The authors gratefully acknowledge the financial support of the Belgian National Fund for Scientific Research - FNRS and Leandro M. Alonso for his python codes used for the currentscape generation.

## Methods

All simulations were performed using the Julia programming language. Analysis were performed either in Matlab and Excel. Code files are freely available at https://github.com/KJacquerie/codes.

### Conductance-based modeling

Single-compartment Hodgkin-Huxley models are used for all neuron models where the membrane potential,*V_m_*, evolves as follows:

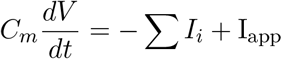

where *C_m_* is the membrane capacitance, *I_i_* is a current due to ionic channels, I_app_ is an external applied current. Ionic currents are voltage-dependent; they are expressed as follows:

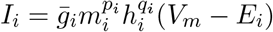

where 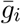 is the maximal conductance, *m_i_* is the activation variable, *h_i_* is the inactivation variable, *p_i_* is an integer between 1 and 4, *q_i_* is either 0 or 1, and *E_i_* i the reversal potential of the channel. The activation and inactivation variable evolve such as:

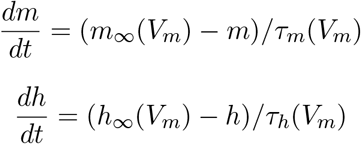

where *m_∞_* and *h_∞_* are the steady-state values of the activation and inactivation variables, and *τ_m_* and *τ_h_* are their respective voltage-dependent time constants. Functions and parameter values are listed in Appendix for each model.

The computational models are run on Julia, a programming language. The ordinary differential equations (ode) are solved with an Euler explicit method. The step time is adapted depending on the model.

### Computational experiment at single-cell

Figure 1A is generated with the model described in [Drion et al. (2018)]. The external current, which hyperpolarizes the cell after 0.5s, switches the firing activity from tonic mode to bursting mode.

Figure 1B is constructed following the method described in [Alonso and Marder (2019)]. The currentscape helps to visualize the contribution of each ionic current as the percentage of the total current.

Figure 1C displays traces of models 1, 5 and 5’ simulated for two values of the membrane capacitance: the nominal value *C_m_* and when it is divided by 10 *C_m_/*10. The current protocol is depolarizing during 0.5s and then hyperpolarizing during 1.5s.

Figure 1D is a quantitative representation of the model capability to switch from tonic to burst for several values of the membrane capacitance *C_m_*. A depolarizing current is applied during 1.5s followed by a hyperpolarizing current during 5.5s. We automatically check the rhythmic pattern; we wait for 0.5s to take into account transient effect, then we record spike times. A spike is considered when the voltage value is greater than −10mV and we save the time at which the event occurs. From spike times, we can deduce the firing pattern. A cell that has no recorded spike time is defined as *silent*. The distinction between tonic mode and bursting mode is based on the comparison of the maximal and minimal interspike-interval (ISI). A neuron is *bursting* when the maximal value of ISI is three times greater than the smallest ISI (max[ISI] > 3 min[ISI]). It ensures the cell to have clusters of action potentials separated from each other by silent intervals. If this criterion is not respected, the cell is classified as a *tonically firing* cell with regular action potential generations [Drion et al. (2018); Goldman et al. (2000)]. For the nominal parameter set, the value of I_app_ is known to generate tonic mode. Afterwards, the membrane capacitance *C_m_* is altered with a multiplicative factor varying from 0.01 to 0.1 with a step of 0.01 ([0.01:0.01:0.1]*C_m_*), then from 0.1 to 5 with a step of 0.1 ([0.1:0.1:5] *C_m_*). In addition, at each tested value of *C_m_*, we scan several values I_app_ in order to find the largest range of capacitance values leading to a switch from tonic to burst. The step time is also adapted for small values of the membrane capacitance to guarantee stability when solving the ode with Euler’s method (for numerical values see SI).

### Computational experiment on a 2-cell circuit

Figure 2A (left) illustrates a 2-cell circuit in a typical thalamic configuration. Neuron models were connected via AMPA, GABA_A_ and GABA_B_ synapses. AMPA synapse provides an excitatory current while GABA is used as an inhibitory current. They are modeled as follows:

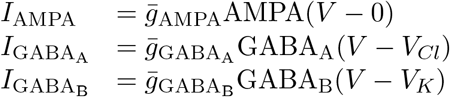

where 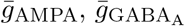 and 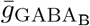 are the synaptic weights. AMPA receptor reversal potential is set to 0mV, GABA_A_ receptor reversal potential is set to chloride reversal potential (*V_Cl_* = − 70mV) and GABA_B_ receptor reversal potential is set to potassium reversal potential (*V_K_* = − 85mV). AMPA, GABA_A_ and GABA_B_ are variables whose dynamics depends on the presynaptic potential *V_pre_* following the equations

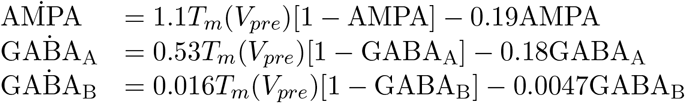

with 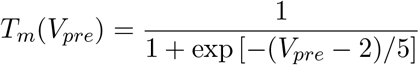

Synaptic weights (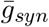 referring to 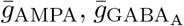 and 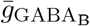) are initially chosen for each model in a way that the 2-cell circuit performs a rhythmic transition when the inhibitory cell is hyperpolarized. Figure 2A, center panel illustrates traces of the desired outcome (recording from model 1 with two identical cells). By contrast, Figure 2A, right panel exhibits an arrhythmic state (recording from model 1 when 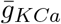 is divided by 10 and 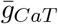 is divided by 5).

Figure 2B is a quantitative comparison of model capability to switch for an increasing intrinsic variability. The intrinsic feature refers to maximal ionic conductances 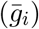. Each maximal conductance is randomly picked with respect to a uniform distribution in a fixed interval around its nominal value. The interval width defines the variability level, as a percentage around this nominal value. For instance, for an intrinsic variability of 10%, each ionic conductance is selected in the range: 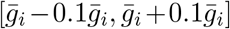. Regarding synaptic connections, each synaptic weight 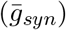 is taken randomly with respect to uniform distribution around its nominal value in this interval 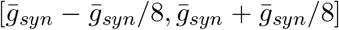. For each model, one thousand sets of parameters are generated in order to build one thousand 2-cell circuits.

Each circuit is simulated during 82s. The external current depolarizes the inhibitory cell during 41s and then hyperpolarizes it during 41s. After 1s of transient period in each state, we record the spike time, *i.e.* a spike is considered when the membrane voltage is greater than −20mV. Then we identify the firing pattern of each cell based on its spike times. A cell is silent when it has not fired. A cell is bursting when the maximal interspike interval is four times greater than the minimum interspike interval, otherwise, the cell is in tonic mode. Then, we identify if the circuit is performing a rhythmic transition. During the depolarized state, the excitatory cell is silent and the inhibitory cell is spiking. During the hyperpolarized state, both cells are synchronously bursting.

### Computational experiment on a 2-cell circuit with a varying the T-type calcium activation time constant

Figure 3A shows the excitatory-inhibitory 2-cell circuit affected by neuromodulation, synaptic plasticity and a tunable time constant for the T-type calcium channel activation 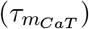. For each model, we generate 400 different 2-cell circuits. For each circuit, maximal intrinsic conductances and synaptic conductances are randomly picked with respect to a uniform distribution in an interval of 20% around their nominal value 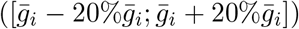. These 400 circuits associated with 400 sets of conductances are simulated for varying CaT channel activation time constant; 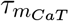 is scaled with the following multiplicative factors; [1/100, 1/50, 1/20, 1/10, 1/8, 1/5, 1/4, 1/3, 1/2, 1/1.5, 1, 1.5, 2, 3, 4, 5, 8, 10, 20, 50, 100]. For each model, we check automatically, at every scaled 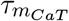, how many circuits among the 400 simulated have performed a rhythmic transition according the same procedure as described above. Figure 3B summarizes the numerical values of 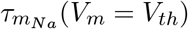, 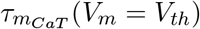 and 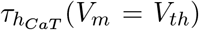. Since the time constants are voltage-dependent, to compare them we fix *V_m_* equal to a threshold voltage, *V_th_*. The threshold voltage for calcium channel activation is chosen at the beginning of the spike upstroke.The threshold voltage for sodium channel activation is chosen at the spike initiation depending on each model. The threshold voltage for calcium channel inactivation is fixed at the beginning of the calcium spike (see SI for numerical values). On Figure 3C, the x-axis represents the *scaled* CaT activation time constant *i.e.* the multiplicative factor times 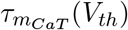 displayed on a logarithmic scale. In order to graduate the axis with a numerical value, the expression is evaluated at the threshold voltage *V_m_* = *V_th_*. The y-axis represents the percentage of rhythmic circuits. The time constants of the Na channel activation, 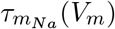 and inactivation 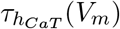 are also marked on the graph. They are evaluated at the threshold voltage *V_m_* = *V_th_*. The results are robust to the choice of the threshold voltage. Models 5’ and 6’ have an instantaneous sodium activation channel, therefore the right boundary is replaced by 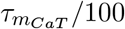.

Figure 3D combines the individual model robustness analysis on one graph. Since channel activation or inactivation time constants have not the same order of magnitude between each model, the x-axis is a *normalized* logarithmic scale. It is normalized such as the left (resp. right) boundary is the time constant of the Na channel activation (resp. CaT channel inactivation) evaluated at its threshold voltage. We scale the axis with a logarithmic scale such as 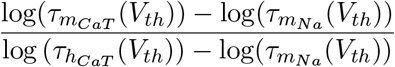 where 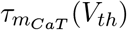 is affected by the multiplicative factor. The idea was to superpose every model between their respective fast timescale (associated with 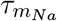) and ultraslow timescale (associated with 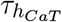). It corresponds to scale each individual plot from Figure 3C in the window bounded by 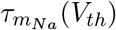 and 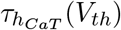

### Computational experiment on a 200-cell network

Figure 4A (resp. C) displays the network configuration and the connectivity between the excitatory and inhibitory neuron populations for a homogeneous (resp. heterogeneous) network. E-cells are connected to I-cells via AMPA synapses and I-cells are connected to E-cells via GABA_A_ and GABA_B_. A homogeneous network is built with 200 identical neurons, and the synaptic weights are the same between each neuron. By contrast, a heterogeneous network is built with neuron models whose maximal ionic conductances are randomly picked in an interval whose width is model-dependent. The interval width is equal to 20% around their nominal values for models 1,2,3,5’, 10% for models 4 and 6’ and 5% for model 5. Synaptic weights are taken randomly with respect to uniform distribution around its nominal value in this interval 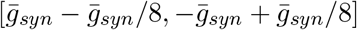.

Analyzing activity in a large neuron population is performed through the computation of the local field potential (LFP). It is the mean field measure of the average behavior of interacting neurons [Buzsáki (2009); Destexhe (1998); Lee and Dan (2012)]. Figures 4C and D (top curves) reflect the collective synaptic activity of the neuronal population. It is modeled by the normalized sum of the post-synaptic currents: 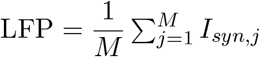 where *M* is the number of post-synaptic neurons in the population. The post-synaptic current from neuron *i* to neuron *j* is

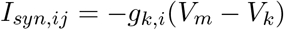

where *k* is the receptor type (AMPA, GABA_A_ and GABA_B_). The entire post synaptic current of the neuron *j* is the sum of the post-synaptic current for all the neighboring neurons. The sum over the *N* neuron is computed:

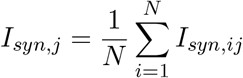

where *N* is the number of pre-synaptic neurons to the neuron *j*. Finally, to compute the local field potential, the sum of all the post-synaptic current of all the neurons are given by:

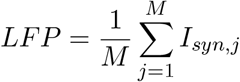

where *M* is the number of post-synaptic neurons in the population.

The LFPs are low-pass filtered at 100Hz via a fourth order Butterworth filter reflecting the use of macro-electrodes in LFP acquisition. The time-frequency plot shows the evolution of the frequency content along time. It results from a logarithmic representation of the spectrogram LFP obtained via a short-time Fourier transform [Drion et al. (2018)]. Spectrograms are obtained via Matlab function spectrogram and the input parameters such as the sampling frequency, the time window, the overlap period, the signal-noise-ration (SNR) are adapted for each model (see SI for numerical values).

Figures 4B and D illustrate the spectrogram of the LFP of the inhibitory neuron population for model 1 (resp. model 5) on the left (resp. right) panel. The homogeneous and the heterogeneous network are respectively on top and bottom. The simulation is performed during 42s split into 21s of depolarized state followed by 21s of hyperpolarized state. When the network is in oscillatory state, the spectrogram is marked by a high power LFP frequency band (yellow band) characterizing that the mean-field rhythmic activity is turned on.

### Computational experiment on a 200-cell network with a varying T-type calcium activation time constant

Figure 5 reproduces the 200-cell network of the Figure 4 with the same current protocol. The 200-cell network is built by randomly picked intrinsic and synaptic conductances in a given interval around their nominal values. The width of the interval is successively increased with steps of 5% around the nominal values (from 0% to 50%). At each variability order, the heterogeneous network sees the CaT activation time constant 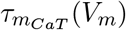 of every neuron scaled by a multiplicative factor equal to [1/8, 1/5, 1/4, 1/3, 1/2, 1, 2, 3, 4, 5, 8]. The LFP activity of the neuron population is plotted similarly as in Figure 4. At each scaled time constant, we detect if the mean-field activity is turned on by analyzing the time course and the spectrogram of the LFP. If the time-course shows a transition in active state to an oscillatory state and if the spectrogram is marked by a high power band during the hyperpolarized state, the network is considered to switch. We summarize the computational experiment in a table showing vertically the different multiplicative factor and the color shows the largest width of the variability interval at which the network switches.

