## Supplementary material for "Robust switches in thalamic network activity require a timescale separation between sodium and T-type calcium channel activations": Jacquerie2020_SI.pdf

Kathleen Jacquerie and Guillaume Drion
Department of Electrical Engineering and Computer Science, University of Liege, Belgium

### Supplementary Information

All simulations were performed using the Julia programming language. Analysis were performed either in Matlab and Excel . Code files are freely available at `https://github.com/KJacquerie/codes`.

#### Model 1

*Paper:* [Drion, Dethier, Franci, and Sepulchre (2018)]

*Ionic currents*
This model is composed of a six ionic currents:

- a transient sodium current: $I_{Na} = \bar{g}_{Na} m_{Na}^3 h_{Na} (V_m - V_{Na})$,
- a delayed-rectifier potassium current: $I_{K,D} = \bar{g}_{K,D} m_{K,D}^4 (V_m - V_K)$,
- a T-type calcium current: $I_{CaT} = \bar{g}_{CaT} m_{CaT}^3 h_{CaT} (V_m - V_{Ca})$,
- a calcium-activated potassium current: $I_{K,Ca} = \bar{g}_{K,Ca} m_{K,Ca\infty}(Ca)(V_m - V_K)$,
- a hyperpolarization-activation cation current: $I_H = \bar{g}_H m_H (V_m - V_H)$,
- a leak current: $I_{leak} = \bar{g}_{leak}(V_m - V_{leak})$.

Voltage-dependent steady-state functions and associated time constants for activation and inactivation gating variables are given in Table 2. The calcium-dependent activation of the calcium-activated potassium current is modeled as follows: $m_{K,Ca,\infty}([Ca]) = ([Ca]/([Ca] + K_D))^2$. The calcium dynamics follows the equation $[\dot{Ca}] = -k_1 I_{CaT} - k_2 [Ca]$ where $k_1$ and $k_2$ are rate parameters. Parameters used in simulations are: $C_m = 1[\mu F/cm^2]$, $V_{Na} = 50$mV, $V_K = -85$mV, $V_{Ca} = 120$mV, $V_{leak} = -55$mV, $V_H = -20$mV, $K_D = 170$. The maximal conductances are expressed in $[mS/cm^2]$ and their nominal values are $g_{Na} = 170$, $g_{K,D} = 40$, $g_{CaT} = 0.55$, $g_{K,Ca} = 4$, $g_H = 0.01$ and $g_{leak} = 0.055$

*Connectivity*
For circuit and network models, the nominal values of maximal synaptic conductances are equal to: $\bar{g}_{\mathrm{AMPA}} = 0.1/n_E$, $\bar{g}_{\mathrm{GABA_A}} = 0.4/n_I$ and $\bar{g}_{\mathrm{GABA_B}} = 2/n_I$ where $n_E$ and $n_I$ are respectively the number of excitatory and inhibitory cells.

*Applied currents*
- At single-cell level, a hyperpolarized-induced bursting is obtained with the applied current equals to 1 then -0.9 $[\mu A/cm^2]$ (referring to Figure 1A).
- Single-cell robustness in $C_m$; the applied current during the first period is equal to 1 and then it is swept from 0 to -2 with a step of 0.1. The hyperpolarized current leading to the highest robustness in capacitance value variation is -1 (referring to Figure 1C).
- For 2-cell circuit or 200-cell network , the applied current exerted on the inhibitory cells is switching

from 1 to -2.6 $[\mu A/cm^2]$ (referring to Figures 2 to 5).

*Threshold voltage*
In Figure 3B, C and D, the voltage-dependent time constants of sodium channel activation $\tau_{m_{Na}}(V_m)$ and T-type calcium channel activation $\tau_{m_{CaT}}(V_m)$ are evaluated at $V_m = -50mV$. The T-type calcium channel inactivation $\tau_{h_{CaT}}(V_m)$ is evaluated at $V_m = -70mV$.

*Currentscapes*
Figure 1B shows the switch from tonic to burst in a neuron model whose parameters are: $C$ =1, $\bar{g}_{Na}$ = 200, $\bar{g}_K$ = 20, $\bar{g}_{CaT}$ = 0.75, $\bar{g}_{KCa}$ = 4, $\bar{g}_H$ = 0.1 (top)/$\bar{g}_H$ = 0.0008 (bottom), $\bar{g}_{leak}$ = 0.055, . The current in the depolarized state is equal to 1 and in the hyperpolarized state -1.85 (top) / -1.07 (bottom). The other parameters remain the same as in the initial description.

#### Model 2

*Paper*: [Destexhe, Contreras, Steriade, Sejnowski, and Huguenard (1996)]

*Ionic currents*
The model is composed of four ionic currents:

- a sodium current: $I_{Na} = \bar{g}_{Na} m_{Na}^3 h_{Na}(V_m - V_{Na})$,
- a potassium current: $I_K = \bar{g}_K m_K^4 (V_m - V_K)$,
- a T-type calcium current: $I_{CaT} = \bar{g}_{CaT} m_{CaT}^2 h_{CaT}(V_m - V_{Ca})$,
- a leakage current: $I_l = \bar{g}_{leak}(V_m - V_{leak})$.

Voltage-dependent steady-state functions and associated time constants for activation and inactivation gating variables are given in Table 3 with $V_2(V) = V - V_{Traub}$. Parameters used in simulations are: $C_m = 1e-3$ $[mF/cm^2]$, $V_{Na} = 50$mV, $V_K = -100$mV, $V_{Ca} = 120$mV, $V_{leak} = -82$mV, $V_{Traub} = -63$mV. Here, we chose to fix the reversal potential of calcium channel rather than evaluate it through calcium concentrations. The maximal conductances are expressed in $[S/cm^2]$ and their nominal values are $\bar{g}_{Na} = 0.4$, $\bar{g}_{K,D} = 0.08$, $\bar{g}_{CaT} = 0.006$, $\bar{g}_{leak} = 5e-5$.

*Connectivity*
For circuit and network models, the nominal values of maximal synaptic conductances are equal to: $\bar{g}_{\mathrm{AMPA}} = 0.1e^{-3}/n_E$, $\bar{g}_{\mathrm{GABA_A}} = 0.2e^{-3}/n_I$ and $\bar{g}_{\mathrm{GABA_B}} = 1e^{-3}/n_I$ where $n_E$ and $n_I$ are respectively the number of excitatory and inhibitory cells.

*Applied currents*
- At single-cell level, a hyperpolarized-induced bursting is obtained with the applied current equals to $0.4e^{-3}$ then 0 $[mA/cm^2]$ (referring to Figure 1A).
- Single-cell robustness in $C_m$; the applied current during the first period is equal to $0.4e^{-3}$ and then it is swept from $0.25e^{-3}$ to -$0.25e^{-3}$ with a step of $0.15e^{-3}$. The hyperpolarized current leading to the highest robustness in capacitance value variation is $0.1e^{-3}$. (referring to Figure 1C).
- For a 2-cell circuit or 200-cell network, the applied current exerted on the inhibitory cells is switching from $0.4e^{-3}$ to -$0.3e^{-3}$ $[mA/cm^2]$ (referring to Figures 2 to 5).

*Threshold voltage*
In Figure 3B, C and D, the voltage-dependent time constants of sodium channel activation $\tau_{m_{Na}}(V_m)$ and T-type calcium channel activation $\tau_{m_{CaT}}(V_m)$ are evaluated at $V_m = -60mV$. The T-type calcium channel inactivation $\tau_{h_{CaT}}(V_m)$ is evaluated at $V_m = -70mV$.

#### Model 3

*Paper:* [Destexhe, Neubig, Ulrich, and Huguenard (1998)]

*Ionic currents*
The model is composed of four ionic currents:

- a sodium current: $I_{Na} = \bar{g}_{Na} m_{Na}^3 h_{Na} (V_m - V_{Na})$,
- a potassium current: $I_K = \bar{g}_K m_K^4 (V_m - V_K)$,
- a T-type calcium current: $I_{CaT} = \bar{g}_{CaT} m_{CaT}^2 h_{CaT} (V_m - V_{Ca})$,
- a leakage current: $I_l = \bar{g}_{leak}(V_m - V_{leak})$.

Voltage-dependent steady-state functions and associated time constants for activation and inactivation gating variables are given in Table 4 with $V_2(V) = V - V_{Traub}$ and a voltage-shift of 3mV for calcium current as explained in [Destexhe et al. (1998)]. Parameters used in simulations are: $C_m = 0.88$ $[\mu F/cm^2]$, $V_{Na} = 50$mV, $V_K = -100$mV, $V_{Ca} = 120$mV, $V_{leak} = -70$mV, $V_{Traub} = -52$mV. Here, we chose to fix the reversal potential of calcium channel instead of computing it through calcium concentration and to use a maximum conductance instead of a maximum permeability. The maximal conductances are expressed in $[mS/cm^2]$ and their nominal values are $\bar{g}_{Na} = 100$, $\bar{g}_{K,D} = 100$, $\bar{g}_{CaT} = 3.3$, $\bar{g}_{leak} = 5e-2$.

*Connectivity*
For circuit and network models, the nominal values of maximal synaptic conductances are equal to: $\bar{g}_{\text{AMPA}} = 0.1/n_E$, $\bar{g}_{\text{GABA}_\text{A}} = 0.2/n_I$ and $\bar{g}_{\text{GABA}_\text{B}} = 1/n_I$ where $n_E$ and $n_I$ are respectively the number of excitatory and inhibitory cells.

*Applied currents*
- At single-cell level, a hyperpolarized-induced bursting is obtained with the applied current equals to 1.5 then -0.7$[\mu A/cm^2]$ (referring to Figure 1A).
- Single-cell robustness in $C_m$; the applied current during the first period is equal to 1.5 and then it is swept from -0.1 to -1 with a step of 0.1. The hyperpolarized current leading to the highest robustness in capacitance value robustness is -0.6 (referring to Figure 1C).
- For 2-cell circuit or 200-cell network, the applied current exerted on the inhibitory cells is switching from 1.5 to -1.7 $[\mu A/cm^2]$ (referring to Figures 2 to 5).

*Threshold voltage*
In Figure 3B, C and D, the voltage-dependent time constants of sodium channel activation $\tau_{m_{Na}}(V_m)$ and T-type calcium channel activation $\tau_{m_{CaT}}(V_m)$ are evaluated at $V_m = -40mV$. The T-type calcium channel inactivation $\tau_{h_{CaT}}(V_m)$ is evaluated at $V_m = -60mV$.

#### Model 4

*Papers:* [Huguenard and McCormick (1992); McCormick and Huguenard (1992)]

*Ionic currents*
The model is composed of eleven ionic currents:

- a transient sodium current: $I_{Na} = \bar{g}_{Na} m_{Na}^3 h_{Na} (V_m - V_{Na})$,
- a persistent sodium current: $I_{Nap} = \bar{g}_{Nap} m_{Nap} (V_m - V_{Na})$,
- a T-type calcium current: $I_{CaT} = \bar{g}_{CaT} m_{CaT}^2 h_{CaT} (V_m - V_{Ca})$,

- a calcium-activated potassium current: $I_C = \bar{g}_C m_C (V_m - V_K)$,
- a low threshold calcium current: $I_L = \bar{g}_L m_L^2 (V_m - V_{Ca})$,
- several potassium current: $I_{K_{2a}} = \bar{g}_{K_{2a}} m_{K_2} h_{K_{2,a}} (V_m - V_K)$, $I_{K_{2b}} = \bar{g}_{K_{2b}} m_{K_2} h_{K_{2,b}} (V_m - V_K)$, $I_A = \bar{g}_A m_A^4 h_A (V_m - V_K)$, $I_{A_2} = \bar{g}_{A_2} m_{A_2}^4 h_{A_2} (V_m - V_K)$
- a sodium leakage current: $I_{Na,leak} = \bar{g}_{Na,leak} (V_m - V_{Na})$,
- a potassium leakage current: $I_{K,leak} = \bar{g}_{K,leak} (V_m - V_K)$.

Voltage-dependent steady-state functions and associated time constants for activation and inactivation gating variables are given in Table 5. Parameters used in simulations are: $C_m = 0.29$ $[nF]$, $V_{Na} = 50$mV, $V_K = -105$mV, $V_{Ca} = 120$mV. Here, we chose to fix the reversal potential of calcium channel instead of computed it through calcium concentration. The maximal conductances are expressed in $[mS]$ and their nominal values are $\bar{g}_{Na} = 12$, $\bar{g}_{Na_p} = 7e-3$, $\bar{g}_A = 20e-3$, $\bar{g}_{A_2} = 15e-3$, $\bar{g}_{CaT} = 1$, $\bar{g}_{K_{2,a}} = 38e-3$, $\bar{g}_{K_{2,b}} = 26e-3$, $\bar{g}_C = 1$, $\bar{g}_L = 0.8$, $\bar{g}_{Na,leak} = 2.65e-3$ and $\bar{g}_{K,leak} = 7e-3$.

*Connectivity*
For circuit and network models, the nominal values of maximal synaptic conductances are equal to: $\bar{g}_{\text{AMPA}} = 0.01/n_E$, $\bar{g}_{\text{GABA}_\text{A}} = 0.04/n_I$ and $\bar{g}_{\text{GABA}_\text{B}} = 0.01/n_I$ where $n_E$ and $n_I$ are respectively the number of excitatory and inhibitory cells.

*Applied currents*
- At single-cell level, a hyperpolarized-induced bursting is obtained with the applied current equals to 1 then 0.1 $[\mu A/cm^2]$ (referring to Figure 1A).
- Single-cell robustness in $C_m$; the applied current during the first period is equal to 1 and then it is swept from 0.4 to -0.2 with a step of 0.1. The hyperpolarized current leading to the highest robustness in capacitance value variation is 0.1 (referring to Figure 1C).
- For 2-cell circuit or 200-cell network, the applied current exerted on the inhibitory cells is switching from 1 to 0 $[\mu A/cm^2]$ (referring to Figures 2 to 5).

*Threshold voltage*
In Figure 3B, C and D, the voltage-dependent time constants of sodium channel activation $\tau_{m_{Na}}(V_m)$ and T-type calcium channel activation $\tau_{m_{CaT}}(V_m)$ are evaluated at $V_m = -50mV$. The T-type calcium channel inactivation $\tau_{h_{CaT}}(V_m)$ is evaluated at $V_m = -60mV$.

#### Model 5

*Paper:* [Wang (1994)]

*Ionic currents*
This model is composed of a six ionic currents:

- a transient sodium current: $I_{Na} = \bar{g}_{Na} m_{Na}^3 (0.85 - m_K)(V_m - V_{Na})$,
- a persistent sodium current: $I_{Nap} = \bar{g}_{Nap} m_{Nap,\infty}^3 (V_m - V_{Na})$,
- a potassium current: $I_K = \bar{g}_K m_K^4 (V_m - V_K)$,
- a T-type calcium current: $I_{CaT} = \bar{g}_{CaT} m_{CaT,\infty}^3 h_{CaT} (V_m - V_{Ca})$,
- a sag current: $I_H = \bar{g}_H m_H^2 (V_m - V_H)$,
- a leak current: $I_{leak} = \bar{g}_{leak} (V_m - V_{leak})$.

Voltage-dependent steady-state functions and associated time constants for activation and inactivation gating variables are given in Table 6. Parameters used in simulations are: $C_m = 1[\mu F/cm^2]$, $V_{Na} = 55$mV, $V_K = -80$mV, $V_{Ca} = 120$mV, $V_{leak} = -70$mV, $V_H = -40$mV, $\sigma_K = 10$, $\sigma_{Na} = 6$, $\sigma_{NaP} = -5$, $\theta_h = -79$ and $k_h = 5$. The maximal conductances are expressed in $[mS/cm^2]$ and their nominal values are $\bar{g}_{Na} = 42$, $\bar{g}_{Nap} = 9$, $\bar{g}_K = 30$, $\bar{g}_{CaT} = 1$, $\bar{g}_H = 0.04$ and $\bar{g}_{leak} = 0.12$

*Connectivity*
For circuit and network models, the nominal values of maximal synaptic conductances are equal to: $\bar{g}_{\text{AMPA}} = 0.1/n_E$, $\bar{g}_{\text{GABA}_\text{A}} = 0.4/n_I$ and $\bar{g}_{\text{GABA}_\text{B}} = 4/n_I$ where $n_E$ and $n_I$ are respectively the number of excitatory and inhibitory cells.

*Applied currents*
- At single-cell level, a hyperpolarized-induced bursting is obtained with the applied current equals to 3 then -1.3 $[\mu A/cm^2]$ (referring to Figure 1A).
- Single-cell robustness in $C_m$; the applied current during the first period is equal to 3 and then it is swept from -0.2 to -2 with a step of 0.2. The hyperpolarized current leading to the highest robustness in capacitance value robustness is -1.5 (referring to Figure 1C).
-For 2-cell circuit or 200-cell network, the applied current exerted on the inhibitory cells is switching from 3 to -1.3 $[\mu A/cm^2]$(referring to Figures 2 to 5).

#### Model 5′

The model designed by Wang in 1994 [Wang (1994)] has set the activation variable of the calcium channel as its steady-state value. It means this activation happens instantaneously without any dynamics *i.e.* without any time constant. We build a new version of Wang model and call it WangCa or model 5′. We restore the slow activation of this channel by integrating the initial expression of the channel as described in the Wang model designed in 1991 with $V_s = 2$. The time constant is equal to:

$$\tau_{m_{CaT}} = 2.5\frac{1.7 + \exp\left[\frac{-(V+V_s+28.8)}{13.5}\right]}{1 + \exp\left[\frac{-(V+V_s+63)}{7.8}\right]}$$

The expression is slightly scaled compared to the original one in [Wang, Rinzel, and Rogawski (1991)]. The time constant of the calcium channel inactivation is also scaled with a factor 5 ($\tau_{h_{CaT}}[WangCa] = 5\tau_{h_{CaT}}[Wang]$). The other parameters remain the same except in Figure 1C, the hyperpolarizing current leading the highest robustness in capacitance value variation is equal to -1.9 (instead of -1.5).

*Threshold voltage*
In Figure 3B, C and D, the voltage-dependent time constants of sodium channel activation $\tau_{m_{Na}}(V_m)$ and T-type calcium channel activation $\tau_{m_{CaT}}(V_m)$ are evaluated at $V_m = -40mV$. The T-type calcium channel inactivation $\tau_{h_{CaT}}(V_m)$ is evaluated at $V_m = -60mV$.

#### Model 6

*Paper:* [Rush and Rinzel (1994)]

*Ionic currents*
This model is composed of a five ionic currents:

- a transient sodium current: $I_{Na} = \bar{g}_{Na} m^3_{Na,\infty}(V_m)(0.85 - m_K)(V_m - V_{Na})$,

- a potassium current: $I_K = \bar{g}_K m_K^4 (V_m - V_K)$,
- a T-type calcium current: $I_{CaT} = \bar{g}_{CaT} m_{CaT,\infty}^3 h_{CaT} (V_m - V_{Ca})$,
- a sodium leak current: $I_{Naleak} = \bar{g}_{Naleak}(V_m - V_{Na})$,
- a sodium leak current: $I_{Kleak} = \bar{g}_{Kleak}(V_m - V_K)$.

Voltage-dependent steady-state functions and associated time constants for activation and inactivation gating variables are given in Table 7. Parameters used in simulations are:$C_m = 1[\mu F/cm^2]$, $V_{Na} = 50$mV, $V_K = -85$mV, $V_{Ca} = 120$mV, $\theta_s = -63$, $k_s = -7.8$, $\theta_h = -72$, $k_h = 1.1$,$\sigma_m = 10.3$, $\sigma_n = 9.3$ and $\phi = 1$. The maximal conductances are expressed in $[mS/cm^2]$ and their nominal values are $\bar{g}_{Na} = 120$, $\bar{g}_K = 10$, $\bar{g}_{CaT} = 0.3$, $\bar{g}_{Naleak} = 0.01429$ and $\bar{g}_{Kleak} = 0.08571$

*Connectivity*
For circuit and network models, the nominal values of maximal synaptic conductances are equal to: $\bar{g}_{\mathrm{AMPA}} = 0.1/n_E$, $\bar{g}_{\mathrm{GABA_A}} = 0.4/n_I$ and $\bar{g}_{\mathrm{GABA_B}} = 2/n_I$ where $n_E$ and $n_I$ are respectively the number of excitatory and inhibitory cells.

*Applied currents*
- At single-cell level, a hyperpolarized-induced bursting is obtained with the applied current equals to 15 then -1.2 $[\mu A/cm^2]$ (referring to Figure 1A).
- Single-cell robustness in $C_m$; the applied current during the first period is equal to 15 and then it is swept from -0.6 to -1.3 with a step of 0.1. The hyperpolarized current leading to the highest robustness in capacitance value variation is -1.2 (referring to Figure 1C).
- For 2-cell circuit or 200-cell network, the applied current exerted on the inhibitory cells is switching from 15 to -1.2 $[\mu A/cm^2]$ (referring to Figures 2 to 5).

#### Model 6′

We built is a modified version of the neuron model established by Rush and Rinzel 1994 [Rush and Rinzel (1994)]. While the activation of calcium current was considered as an instantaneous event in [Rush and Rinzel (1994)], we restore the slow activation of the T-type calcium channel. We call this new model RushCa or model 6′. To do so, we added a voltage-dependent function for the time constant of the T-type calcium channel activation and remove the simplification that was performed in the original paper. Therefore, the ionic currents are the same as the initial model except for the T-type calcium current: $I_{CaT} = \bar{g}_{CaT} m_{CaT}^3 h_{CaT}(V_m - V_{Ca})$. The associated time constant is expressed as follows:

$$\tau_{m_{CaT}} = 0.1 \frac{1.7 + \exp\left[\frac{-(V+28.8)}{13.5}\right]}{1 + \exp\left[\frac{-(V+63)}{7.8}\right]}$$

The membrane capacitance is reduced by a factor of 10 such as $C_m = 0.1[\mu F/cm^2]$. In order to respect the physiological timescale of the different ionic currents (such as fast activation and slow inactivation of sodium channels, slow activation of potassium channels, slow activation and ultraslow inactivation of calcium channels), we adapted the time constant of the calcium channel inactivation and the potassium channel activation:$\tau_{h_{CaT}}$[RushCa] $= 1.5\tau_{h_{CaT}}$[Rush] and $\tau_{m_K}$[RushCa]$= 0.175\tau_{m_K}$[Rush]. The synaptic conductances remain the same. The external current is also the same for every simulation. Except the hyperpolarized current leading to the highest robustness in capacitance value variation is -1.1 (instead of -1.2).

*Threshold voltage*
In Figure 3B, C and D, the voltage-dependent time constants of sodium channel activation $\tau_{m_{Na}}(V_m)$

and T-type calcium channel activation $\tau_{m_{CaT}}(V_m)$ are evaluated at $V_m = -40mV$. The T-type calcium channel inactivation $\tau_{h_{CaT}}(V_m)$ is evaluated at $V_m = -50mV$.

#### Model summary

| Model #<br>Model name | 1<br>Drion | 2<br>Destexhe | 3<br>Destexhe 1998 | 4<br>Huguenard and McCormick | 5<br>Wang | 6<br>Rush | $5'$<br>WangCa | $6'$<br>RushCa |
|---|---|---|---|---|---|---|---|---|
| Publication year | 2018 | 1996 | 1998 | 1992 | 1994 | 1994 | - | - |
| Number of ionic currents | 6 | 4 | 4 | 11 | 6 | 5 | 6 | 5 |
| T-type calcium channel activation dynamics | slow | slow | slow | slow | instant. | instant. | slow | slow |

**Table 1:** Key-information related to the conductance-based models

#### Ionic channel description: steady-state functions and time constants of the gating variables

| $I_i$ | gating variable | time constant |
|---|---|---|
| $I_{Na}$ | $m_{Na,\infty} = \frac{1}{1+\exp[(V+35.5)/-5.29]}$ | $\tau_{m,Na} = 1.32 - \frac{1.26}{1+\exp[(V+120)/-25]}$ |
| | $h_{Na,\infty} = \frac{1}{1+\exp[(V+48.9)/5.18]}$ | $\tau_{h,Na} = \frac{0.67}{1+\exp[(V+62.9)/-10.0]} * \left(1.5 + \frac{1}{1+\exp[(V+34.9)/3.6]}\right)$ |
| $I_{K,D}$ | $m_{K,D,\infty} = \frac{1}{1+\exp[(V+12.3)/-11.8]}$ | $\tau_{m,KD} = 7.2 - \frac{6.4}{1+\exp[(V+28.3)/-19.2]}$ |
| $I_{CaT}$ | $m_{CaT,\infty} = \frac{1}{1+\exp[(V+67.1)/-7.2]}$ | $\tau_{m,CaT} = 21.7 - \frac{21.3}{1+\exp[(V+68.1)/-20.5]}$ |
| | $h_{CaT,\infty} = \frac{1}{1+\exp[(V+80.1)/5.5]}$ | $\tau_{h,CaT} = 410 - \frac{179.6}{1+\exp[(V+55)/-16.9]}$ |
| $I_H$ | $m_{H,\infty} = \frac{1}{1+\exp[(V+80)/6]}$ | $\tau_{m,H} = 272 + \frac{1149}{1+\exp[(V+42.2)/-8.73]}$ |

**Table 2:** Steady-state functions for channel gating variables and time constants for the different ion channels present in Drion model (Model 1).

| $I_i$ | gating variable | time constant |
|---|---|---|
| $I_{Na}$ | $\alpha_{m_{Na}} = \frac{0.32(13-V_2(V))}{\exp\left[\frac{13-V_2(V)}{4}\right]-1}$ | $\tau_{m,Na} = \frac{1}{\alpha_{m_{Na}}+\beta_{m_{Na}}}$ |
| | $\beta_{m_{Na}} = \frac{0.28(V_2(V)-40)}{\exp\left[\frac{V_2(V)-40}{5}\right]-1}$ | |
| | $\alpha_{h_{Na}} = 0.128\exp\left[\frac{17-V_2(V)}{18}\right]$ | $\tau_{h,Na} = \frac{1}{\alpha_{h_{Na}}+\beta_{h_{Na}}}$ |
| | $\beta_{h_{Na}} = \frac{4}{1+exp\left[\frac{40-V_2(V)}{5}\right]}$ | |
| $I_K$ | $\alpha_{m_K} = \frac{0.032(15-V_2(V))}{\exp\left[\frac{15-V_2(V)}{5}\right]-1}$ | $\tau_{m,K} = \frac{1}{\alpha_{m_K}+\beta_{m_K}}$ |
| | $\beta_{m_K} = 0.5\exp\left[\frac{10-V_2(V)}{40}\right]$ | |
| $I_{CaT}$ | $m_{CaT,\infty} = \frac{1}{1+\exp\left[\frac{-(V+50)}{7.4}\right]}$ | $\tau_{m_{CaT}} = 1 + \frac{0.33}{\exp\left[\frac{-(V+100)}{15}\right]+\exp\left[\frac{V+25}{10}\right]}$ |
| | $h_{CaT,\infty} = \frac{1}{1+\exp\left[\frac{V+80}{5}\right]}$ | $\tau_{h_{CaT}} = 28.3 + \frac{0.33}{\exp\left[\frac{V+48}{4}\right]+\exp\left[\frac{-(V+407)}{50}\right]}$ |

**Table 3:** Steady-state functions for channel gating variables and time constants for the different ion channels present in Destexhe,1996 model (Model 2).

| $I_i$ | gating variable | time constant |
| --- | --- | --- |
| $I_{Na}$ | $\alpha_{m_{Na}} = \dfrac{0.32(13 - V_2(V))}{\exp\left[\frac{13-V_2(V)}{4}\right] - 1}$ | $\tau_{m,Na} = \dfrac{1}{\alpha_{m_{Na}} + \beta_{m_{Na}}}$ |
| | $\beta_{m_{Na}} = \dfrac{0.28(V_2(V) - 40)}{\exp\left[\frac{V_2(V)-40}{5}\right] - 1}$ | |
| | $\alpha_{h_{Na}} = 0.128\exp\left[\frac{17-V_2(V)}{18}\right]$ | $\tau_{h,Na} = \dfrac{1}{\alpha_{h_{Na}} + \beta_{h_{Na}}}$ |
| | $\beta_{h_{Na}} = \dfrac{4}{1 + exp\left[\frac{40-V_2(V)}{5}\right]}$ | |
| $I_K$ | $\alpha_{m_K} = \dfrac{0.032(15 - V_2(V))}{\exp\left[\frac{15-V_2(V)}{5}\right] - 1}$ | $\tau_{m,K} = \dfrac{1}{\alpha_{m_K} + \beta_{m_K}}$ |
| | $\beta_{m_K} = 0.5\exp\left[\frac{10-V_2(V)}{40}\right]$ | |
| $I_{CaT}$ | $m_{CaT,\infty} = \dfrac{1}{1 + \exp\left[\frac{-(V+59)}{6.2}\right]}$ | $\tau_{m_{CaT}} = 0.204 + \dfrac{0.333}{\exp\left[\frac{V+18.8}{18.2}\right] + \exp\left[\frac{-(V+134)}{16.7}\right]}$ |
| | $h_{CaT,\infty} = \dfrac{1}{1 + \exp\left[\frac{V+83}{4}\right]}$ | for $V < -80$ : $\tau_{h_{CaT}} = 0.33\exp\left[\frac{V+469}{66.6}\right]$ |
| | | for $V \geq -80$ : $\tau_{h_{CaT}} = 9.32 + 0.33\exp\left[\frac{V+24}{10.5}\right]$ |

**Table 4:** Steady-state functions for channel gating variables and time constants for the different ion channels present in Destexhe,1998 model (Model 3).

| $I_i$ | gating variable | time constant |
|---|---|---|
| $I_{Na}$ | $\alpha_{m_{Na}} = \dfrac{0.091(V+38)}{1-\exp\left[\frac{-(V+38)}{5}\right]}$ | $\tau_{m,Na} = \dfrac{1}{\alpha_{m_{Na}}+\beta_{m_{Na}}}$ |
| | $\beta_{m_{Na}} = \dfrac{-0.062(V+38)}{1-\exp\left[\frac{V+38}{5}\right]}$ | |
| | $\alpha_{h_{Na}} = 0.016\exp\left[\frac{-(V+55)}{15}\right]$ | $\tau_{h,Na} = \dfrac{1}{\alpha_{h_{Na}}+\beta_{h_{Na}}}$ |
| | $\beta_{h_{Na}} = \dfrac{2.07}{\exp\left[\frac{17-V}{21}\right]+1}$ | |
| $I_{Nap}$ | $m_{Nap,\infty} = \dfrac{1}{1+\exp\left[\frac{-(V+49)}{5}\right]}$ | $\alpha_{m_{Nap}} = \dfrac{0.091(V+38)}{1-\exp\left[\frac{-(V+38)}{5}\right]}$, $\beta_{m_{Nap}} = \dfrac{-0.062(V+38)}{1-\exp\left[\frac{V+38}{5}\right]}$ |
| | | $\tau_{m,Nap} = \dfrac{1}{\alpha_{m_{Nap}}+\beta_{m_{Nap}}}$ |
| $I_L$ | $\alpha_{m_L} = \dfrac{1.6}{1+\exp\left[-0.072(V-5)\right]}$ | $\tau_{m,L} = \dfrac{1}{\alpha_{m_L}+\beta_{m_L}}$ |
| | $\beta_{m_L} = \dfrac{0.02(V-1.31)}{\exp\left[\frac{V-1.31}{5.36}\right]-1}$ | |
| $I_C$ | $\alpha_{m_C} = 2.5e5\,[CaL]\exp(V/24)$ | $\tau_{m,C} = \dfrac{1}{\alpha_{m_C}+\beta_{m_C}}$ |
| | $\beta_{m_C} = 0.1\exp[-V/24]$ | |
| $I_{CaT}$ | $m_{CaT,\infty} = \dfrac{1}{1+\exp\left[\frac{-(V+57)}{6.2}\right]}$ | $\tau_{m_{CaT}} = 0.612 + \dfrac{1}{\exp\left[\frac{-(V+131.6)}{16.7}\right]+\exp\left[\frac{V+16.8}{18.2}\right]}$ |
| | $h_{CaT,\infty} = \dfrac{1}{1+\exp\left[\frac{V+81}{4.03}\right]}$ | for $V < -80$: $\tau_{h_{CaT}} = \exp\left[\frac{V+467}{66.6}\right]$ |
| | | for $V \geq -80$: $\tau_{h_{CaT}} = 28+\exp\left[\frac{-(V+21.88)}{10.2}\right]$ |
| $I_A$ | $m_{A,\infty} = \dfrac{1}{1+\exp\left[\frac{-(V+60)}{8.5}\right]}$ | $\tau_{m_A} = 0.37 + \dfrac{1}{\exp\left[\frac{V+35.82}{19.697}\right]+\exp\left[\frac{V+79.69}{-12.7}\right]}$ |
| | $h_{A,\infty} = \dfrac{1}{1+\exp\left[\frac{V+78}{6}\right]}$ | for $V < -63$: $\tau_{h_A} = \dfrac{1}{\exp\left[\frac{V+46.05}{5}\right]+\exp\left[\frac{V+238.4}{-37.45}\right]}$ |
| | | for $\tau_{h_A} = 19$ |
| $I_{A,2}$ | $m_{A2,\infty} = \dfrac{1}{1+\exp\left[\frac{-(V+36)}{20}\right]}$ | $\tau_{m_{A2}} = 0.37 + \dfrac{1}{\exp\left[\frac{V+35.82}{19.697}\right]+\exp\left[\frac{V+79.69}{-12.7}\right]}$ |
| | $h_{A2,\infty} = \dfrac{1}{1+exp\left[\frac{V+78}{6}\right]}$ | for $V < -73$: $\tau_{h_{A2}} = \dfrac{1}{\exp\left[\frac{V+46.05}{5}\right]+\exp\left[\frac{V+238.4}{-37.45}\right]}$ |
| | | for $V \geq -73$ $\tau_{h_{A2}} = 60$ |
| $I_{K2}$ | $m_{K2,\infty} = \dfrac{1}{1+\exp\left[\frac{V+43}{-17}\right]}$ | $\tau_{m_{K2}} = 9.9 + \dfrac{1}{\exp\left[\frac{V-81}{25.6}\right]+\exp\left[\frac{V+132}{-18}\right]}$ |
| | $h_{K2a,\infty} = \dfrac{1}{1+\exp\left[\frac{V+58}{10.6}\right]}$ | $\tau_{h_{K2a}} = 120 + \dfrac{1}{\exp\left[\frac{V-1.329}{200}\right]+\exp\left[\frac{V+130}{-7.1}\right]}$ |
| $I_{K2b}$ | $h_{K2b,\infty} = \dfrac{1}{1+\exp\left[\frac{V+58}{10.6}\right]}$ | for $V < -70$: $\tau_{h_{K2b}} = 120 + \dfrac{1}{\exp\left[\frac{V-1.329}{200}\right]+\exp\left[\frac{V+130}{-7.1}\right]}$ |
| | | for $V \geq -70$: $\tau_{h_{K2b}} = 8.9$ |

**Table 5:** Steady-state functions for channel gating variables and time constants for the different ion channels present in Huguenard and McCormick model (model 4).

| $I_i$ | gating variable | time constant |
|---|---|---|
| $I_{Na}$ | $\alpha_{m_{Na}} = \dfrac{-0.1(V+29.7-\sigma_{Na})}{\exp\left[-0.1(V+29.7-\sigma_{Na})\right]-1}$ | - |
| | $\beta_{m_{Na}} = 4\exp\left[\frac{-(V+54.7-\sigma_{Na})}{18}\right]$ | |
| $I_{Nap}$ | $\alpha_{m_{Nap}} = \dfrac{-0.1(V+29.7-\sigma_{Nap})}{\exp\left[-0.1(V+29.7-\sigma_{Nap})\right]-1}$ | - |
| | $\beta_{m_{Nap}} = 4\exp\left[\frac{-(V+54.7-\sigma_{Nao})}{18}\right]$ | |
| $I_K$ | $\alpha_{m_K} = \dfrac{-0.01(V+45.7-\sigma_K)}{\exp\left[-0.1(V+45.7-\sigma_K)\right]-1}$ | $\tau_{m,K} = \dfrac{7/200}{\alpha_{m_K}+\beta_{m_K}}$ |
| | $\beta_{m_K} = 0.125\exp\left[\frac{-(V+55.7-\sigma_K))}{80}\right]$ | |
| $I_{CaT}$ | $m_{CaT,\infty} = \dfrac{1}{\exp\left[\frac{-(V+65)}{7.8}\right]}$ | - |
| | $h_{CaT,\infty} = \dfrac{1}{\exp\left[\frac{(V-\theta_h)}{k_h}\right]}$ | $\tau_{h,CaT} = \frac{1}{2}\left\{\dfrac{\exp\left[\frac{V+162.3}{17.8}\right]}{\exp\left[\frac{(V-\theta_h)}{k_h}\right]}+20\right\}$ |
| $I_H$ | $m_{H,\infty} = \dfrac{1}{1+\exp\left[\frac{V+69}{7.1}\right]}$ | $\tau_{m_H} = \dfrac{1000}{\exp\left[\frac{V+66.4}{9.3}\right]+\exp\left[\frac{-(V+81.6)}{13}\right]}$ |

**Table 6:** Steady-state functions for channel gating variables and time constants for the different ion channels present in Wang model (Model 5).

| $I_i$ | gating variable | time constant |
|---|---|---|
| $I_{Na}$ | $\alpha_{m_{Na}} = \dfrac{0.1(V+35-\theta_m)}{1-\exp\left[-0.1(V+35-\sigma_m)\right]}$ | - |
| | $\beta_{m_{Na}} = 4\exp\left[-0.05(V+60-\sigma_m)\right]$ | |
| $I_K$ | $\alpha_{m_K} = \dfrac{0.01(V+50-\sigma_n)}{1-\exp\left[-0.1(V+50-\sigma_n)\right]}$ | $\tau_{m,K} = \dfrac{0.05}{\alpha_{m_K}+\beta_{m_K}}$ |
| | $\beta_{m_K} = 0.125\exp\left[-0.0125(V+60-\sigma_n)\right]$ | |
| $I_{CaT}$ | $m_{CaT,\infty} = \dfrac{1}{\exp\left[\frac{V-\theta_s}{ks}\right]}$ | - |
| | $h_{CaT,\infty} = \dfrac{1}{0.5+\sqrt{0.25+\exp\left[\frac{V-\theta_h}{kh}\right]}}$ | $\tau_{h,CaT} = 1/\phi\left\{\dfrac{\exp\left[\frac{V+150}{18}\right]}{1.5+\sqrt{0.25+\exp\left[\frac{V-80}{4}\right]}}+30\right\}$ |

**Table 7:** Steady-state functions for channel gating variables and time constants for the different ion channels present in Rush model (Model 6).
